# Endonucleolytic RNA cleavage drives changes in gene expression during the innate immune response

**DOI:** 10.1101/2023.09.01.555507

**Authors:** Agnes Karasik, Hernan A. Lorenzi, Andrew V. DePass, Nicholas R. Guydosh

## Abstract

Viral infection triggers several dsRNA sensors that lead to changes in gene expression in the cell. One of these sensors activates an endonuclease, RNase L, that cleaves single stranded RNA. However, how the resultant widespread RNA fragmentation affects gene expression is not fully understood. Here we show that this fragmentation induces the Ribotoxic Stress Response via ZAKα, potentially through ribosome collisions. The p38 and JNK pathways that are activated as part of this response promote outcomes that inhibit the virus, such as programmed cell death. We also show that RNase L limits the translation of stress-responsive genes, including antiviral *IFIT* mRNAs and *GADD34* that encodes an antagonist of the Integrated Stress Response. Intriguingly, we found the activity of the generic endonuclease, RNase A, recapitulates many of the same molecular phenotypes as activated RNase L, demonstrating how widespread RNA cleavage can evoke an antiviral program.

**Highlights:**

- Activated RNase L acts with dsRNA-sensing pathways to promote cell signaling
- RNA fragmentation induces transcription through ZAKα signaling
- Activation of RNase L modulates levels of eIF2α phosphorylation
- Translation of the *GADD34* and *IFIT* mRNAs is inhibited by active RNase L

## Introduction

Viral infections are recognized by <u>P</u>attern <u>R</u>ecognition <u>R</u>eceptors (PRRs) that trigger the innate immune response and ultimately promote elimination of pathogens. Foreign nucleic acid (RNA or DNA) sensing is one of the major ways viral pathogens are recognized ^1^. Viruses have either an RNA genome or produce RNA intermediates, and thus sensing of viral RNA in the cytoplasm is vital for recognizing and fighting against viruses. A subset of PRRs recognize double stranded RNA (dsRNA) ^1^. One group within these PRRs is the interferon-induced family of <u>O</u>ligo-<u>A</u>denylate <u>S</u>ynthetases (OAS1-3) that produce an oligomer of adenylates chained together through 2’-5’ linkages (2-5A) upon dsRNA binding. 2-5A can selectively bind Ribonuclease L (RNase L) to induce dimerization that is required for its transition from a latent to an activated state. Activated RNase L then cleaves a variety of host and viral RNAs, triggering many downstream processes and physiological changes in the cell. These changes include the loss of most mRNAs (and therefore gene expression in the cell), production of chemokines and cytokines, activation of inflammasomes, inhibition of cell migration, autophagy, senescence, and cell death ^2–5^. Intriguingly, the function of RNase L appears to extend beyond its role in protecting against viruses. RNase L mutations increase risk for multiple types of cancer, including prostate and breast cancer ^6–9^, and autoimmune diseases ^10^.

Besides RNase L, another interferon-regulated PRR, <u>P</u>rotein <u>K</u>inase <u>R</u> (PKR), can induce reduction in protein synthesis in the cell through phosphorylation of <u>e</u>ukaryotic Initiation <u>F</u>actor 2α (eIF2α). Phospho-eIF2α inhibits initiation of translation of the majority of mRNAs, presumably to limit viral protein production, and triggers a number of changes in the cell that are collectively referred to as the Integrated <u>S</u>tress <u>R</u>esponse (ISR) ^11–13^. Additionally, the dsRNA receptors RIG-I and MDA5 induce the interferon and inflammatory response by activating signaling that leads to transcription. Activation of the different dsRNA response pathways in the cytoplasm can have intersecting functions, wherby activated RIG-I induces gene expression, while activated RNase L and PKR reduce gene expression. However, how the different dsRNA sensing pathways, particularly RNase L, act together to achieve a strong antiviral response is largely unknown.

Activation of RNase L can induce and alter transcription ^14–17^. During RNase L activation, many proinflammatory genes and cytokines (e.g. CXCL8) were found to be upregulated in early microarray studies ^14^. Importantly, more recent RNA-sequencing (RNA-seq) studies found that most transcripts are degraded (∼60 to >99%) by RNase L. Interestingly, immune transcripts, such as interferon-β (IFNβ), are able to compensate for this degradation via transcriptional upregulation or resistance to cleavage due to underrepresentation of favorable RNase L cleavage motifs (UU or UA) ^16,17^. Furthermore, many Interferon <u>S</u>timulated <u>G</u>enes (ISGs) are upregulated during the dsRNA response in both WT and *RNASEL* KO cells, suggesting that they do not depend on RNase L, whereas upregulation of cytokines, such as CXCL8, CXCL2, and other immune genes, including *IFNL1* and *2*, and *IFIT3*, were at least partially dependent on RNase L ^17^. These observations suggest that active RNase L together with the other dsRNA pathways may amplify transcription of antiviral genes. In some cell lines, RNase-L cleaved RNA fragments play a role in activating RIG-I ^18,19^ and the inflammasome ^20^, however interferon stimulation (downstream of RIG-I) was not observed in human lung carcinoma cells ^16,17,21^. One limitation of the existing studies that used RNA-seq is that RNase L was activated by a dsRNA mimic (poly I:C), an approach that simultaneously activates many other pathways ^16,17^. These pathways can mask RNase L-specific effects and make it difficult to assess how RNase L activation and the other dsRNA pathways act together to induce the antiviral immune response.

Importantly, activation of RNase L can also induce pathways related to the <u>R</u>ibotoxic <u>S</u>tress <u>R</u>esponse (RSR) ^22,23^, which triggers apoptosis through transcription activated by JNK (c-<u>J</u>UN <u>N</u>-terminal <u>K</u>inase) signaling, which is a mitogen-activated protein kinase (MAPK) family member. While several MAP2K family members can phosphorylate JNK ^23^, the mechanism of how they are activated by RNase L has remained unclear. The RSR was originally shown to be activated by defective translation, so it was suggested that pathways related to the RSR were induced when RNase L damaged ribosomes by cleaving rRNA ^22–24^. However, more recently, it was revealed that ribosomes with cleaved rRNAs are functional, suggesting that ribosome damage may not be responsible for activating the RSR ^16^. In addition, the RSR was recently shown to include activation of the MAP3K family member ZAKα that senses collisions between ribosomes and phosphorylates MAP2Ks that, in turn, phosphorylate JNK, and another MAPK, p38 ^25–28^. These observations suggest that activation of ZAKα by ribosome collisions could occur when RNase L induces the RSR. However, the source of any potential ribosome collisions in RNase L activated cells remains unknown. One possible source of ribosome collisions was suggested by our previous observation that activation of RNase L leads to translation of fragmented mRNAs ^21^. Ribosomes that translate mRNA fragments would have the possibility of reaching a 3’ end without terminating, an event known to lead to ribosome stalling and the formation of ribosome collisions ^29^.

More broadly, the overall loss of translated mRNA in the cell upon RNase L activation is expected to change the balance of free ribosomal subunits and mRNAs ^16^. This altered balance could modulate the translation efficiency of the remaining, full-length transcripts. Consistent with this, we previously observed an increase in the relative proportion of ribosomes in alternative <u>O</u>pen <u>R</u>eading <u>F</u>rames (ORFs) or “altORFs” following RNase L activation ^21^. We proposed that translation of these altORFs became more favorable as excess free ribosomes promoted translation initiation at the 5’ ends of the RNA fragments created by RNase L. However, how RNase L and other dsRNA pathways lead to changes in translation on mRNAs remained unclear.

Here we investigated how direct activation of RNase L, and attendant global mRNA degradation, contributes to the activation of the innate immune response. We found that RNase-L mediated RNA cleavage strengthened multiple parts of the antiviral transcriptional response and, in particular, induced ZAKα activation, presumably via ribosome collisions. In addition, activation of RNase L also changed how efficiently intact mRNAs were translated, in some cases by inhibiting the effect of eIF2α phosphorylation. Our results highlight the multifaceted role of RNase L and endonucleolytic cleavage in promoting antiviral functions.

## Results

### Activation of RNase L induces transcription and phosphorylation of ZAKα

To measure changes in the transcriptome under activation of RNase L, but not other components of the dsRNA-induced innate immune response, we performed RNA-seq on wild type (WT) and *RNASEL* KO cells that were transfected with 2-5A (1 µM), the direct and specific activator of RNase L (Figure 1A). Since RNase L cleaves rRNA, we utilized a rRNA degradation assay to assess the level of RNase L activation ^30^. Across all our replicates, we found that rRNA degradation was similar (within a 2-fold difference, Figure S1A). Based on previous reports that used spike-in oligos for normalization ^16,17^, these levels of rRNA degradation correspond to a loss of ∼60% to >99% of individual mRNAs in the cell. We therefore assume that the absolute abundance of most mRNAs in the treated cells under study here substantially decreases (“baseline loss”).

**Figure 1.**
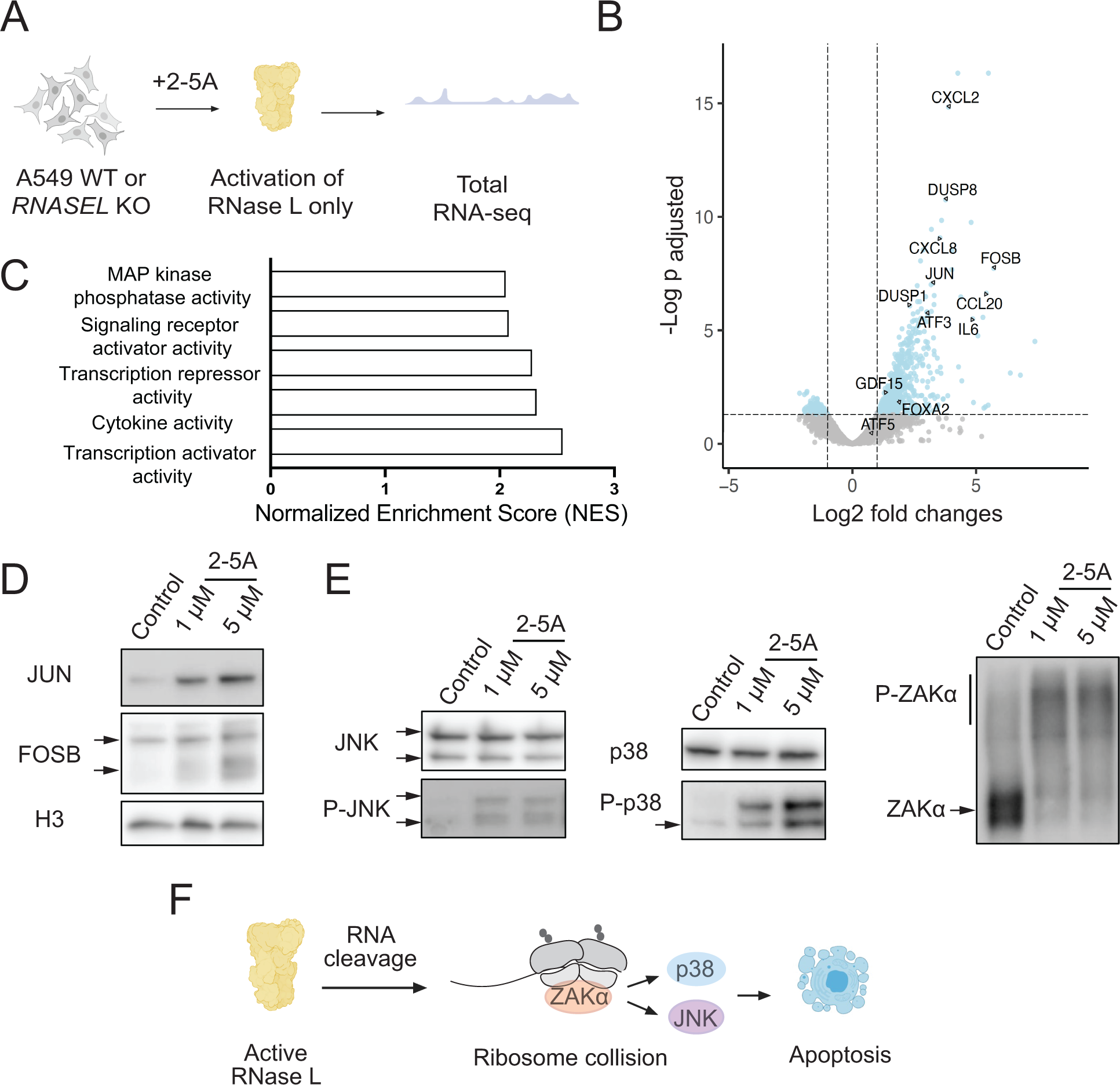
RNase L activation leads to transcriptional changes through ZAK, JNK and p38. **A** Schematics of RNA-seq experiments. A549 cells were transfected with 2-5A and RNA-seq was performed as described in Materials and Methods. **B** Volcano plot showing upregulation of example proinflammatory cytokines and transcription factors during 2-5A treatment. Differentially expressed genes define as p_adjusted_ value <0.05, log_2_fold change >1. **C** Gene Ontology analysis of molecular functions shows that upregulated genes are enriched in genes related to transcription activities. 5 out of 9 categories with the highest Normalized Enrichment Scores are shown where p_adjusted_ <0.05. Categories not shown had highly overlapping gene sets with those that are highlighted. **D** Western blots show that transcription factors FOSB and JUN are increased due to 2-5A activation. Different FOSB isoforms are marked with black arrows. **E** Western blots show JNK and p38 are activated in A549 cells after 2-5A treatment. Different JNK isoforms are marked with black arrows. ZAKα is hyperphosphorylated during RNase L activation. Upper bar shows phosphorylated ZAKα in a Phos-Tag gel (shifted from its expected molecular weight). Lower line points to non-phosphorylated ZAKα. **F** Working model of JNK/p38 pathway activation by RNase L. RNase L cleaves mRNAs that are being translated. At the end of the resulting fragments, ribosomes can collide and activate ZAKα and subsequently JNK and p38.

Then, we analyzed differential expression (DE) patterns in 2-5A treated vs untreated cells by DESeq2 ^31^. The DE changes in our experiments represent differences in relative abundance of the small pool of mRNA that is not degraded. We infer that mRNAs that appear upregulated (described throughout as relative “upregulation”) actually decrease in abundance, but to a lesser extent than the general population, due to resistance to RNase L cleavage activity or compensation by active transcription. The DE values therefore characterize how the global loss of mRNA varies across the transcriptome. We found hundreds of differentially expressed genes (“DE genes,” p_adjusted_<0.05) in 2-5A treated WT cells, revealing that the level of these mRNAs significantly differs from the baseline loss that generally affects all transcripts (Figure 1B). 2-5A treatment did not induce many changes in differential gene expression in *RNASEL* KO cells (Table S1), as expected, since 2-5A is a specific activator of RNase L. DE genes that appeared to be upregulated by RNase L activation (Figure 1B), defined as log2 fold changes >1) included transcripts that encode proinflammatory proteins and cytokines (e.g. *CXCL2*, *CCL20*, *CXCL8*, and *IL6*), suggesting that direct RNase L activation stimulates an inflammatory response. In contrast, we did not detect an interferon (IFN) response, such as increased transcript levels for *IFNB1.* These findings are largely consistent with previous RNA-seq data from human cells (WT vs *RNASEL* KO) that were transfected with poly I:C ^16,17^. These changes were consistent across replicates (Figure S1B).

We noted that some of the genes that appear to be upregulated in 2-5A treated cells encode transcription factors or activators, such as *JUN*, *FOSB*, *FOXA2*, *GDF15*, and *ATF3* (Figure 1B). This is consistent with previous findings for some of these genes, such as *FOXA2* and *GDF15* ^14,32^. Gene Ontology (GO) analysis of molecular functions offered additional support since several of the top categories of DE genes were related to transcription regulation (Figure 1C). This finding suggests that the apparent increase in some transcript levels in our DE analysis could be due to new transcription that is induced by these factors. Since RNase L reduces global mRNA levels in the cell, it is unclear whether the relative upregulation of these transcription factor genes observed in our RNA-seq data is sufficient to increase production of protein. Thus, we probed selected transcription factor protein levels by western blotting. Interestingly, we found that protein levels for FOSB and JUN increased in 2-5A treated WT, but not *RNASEL* KO, cells (Figure 1D and S1C). This finding suggests that activation of RNase L can actually raise the abundance of these proteins, leading to downstream transcriptional activation.

Next, we assessed which pathways are triggered in 2-5A treated cells that could explain the apparent transcriptional upregulation. We found that several target genes in the JNK, p38, ERK, and NF-κB pathways appeared to be increased in our data, suggesting that at least some of these pathways (which are known to have overlapping targets) are activated. For instance, the transcription factor GDF15 controls inflammation, cell repair, and growth, and is known to be upregulated by the activity of the JNK and ERK kinases when prostate cancer cells are treated with 2-5A ^14^. To dissect which of these pathways are specifically responsible for the observed transcriptional changes in our dataset, we used western blotting to assess activation through phosphorylation of these pathways. We found that RNase L can activate JNK and p38 (Figure 1E and S1D), but we did not detect activation of pathways related to ERK (Figure S1E, ERK phosphorylation) or NF-κB (Figure S2, p65 nuclear fraction). These results suggest that transcriptional changes observed in the cells here are driven, in part, by the activation of JNK and p38. We also noticed that multiple *DUSP* mRNAs, such as *DUSP1* and *8*, that encode negative regulators of the JNK pathway were upregulated (Figure 1B and S1B), suggesting a need for modulation of JNK when RNase L is activated. Interestingly, JNK activation can be apoptotic or induce autophagy. Here we observed the induction of proapoptotic genes, such as *JUN*, but not genes expressed during autophagy, such as *ATF5* ^33^.

It has been suggested that activation of RNase L can trigger the RSR, which can lead to downstream activation of the JNK and p38 pathways ^23,25^. Since it was demonstrated that this can involve the ribosome collision sensor and kinase, ZAKα, we tested whether ZAKα is activated in 2-5A treated cells by examining its phosphorylation state by western blotting. We found that ZAKα is strongly activated due to 2-5A treatment in WT but not in *RNASEL* KO cells, indicating that phosphorylation of JNK and p38 under RNase L might occur through ribosome collisions during RNase L activation (Figure 1E-F and S1D).

### The OAS-RNase L pathway acts together with other dsRNA sensing pathways to achieve strong activation of innate immune signaling

We next asked whether activation of RNase L modulates the effects of other pathways that are typically activated during infection by other dsRNA sensors ^18,19^. To simultaneously activate multiple pathways and test RNase L’s role, we transfected WT and *RNASEL* KO A549 cells with poly I:C, a dsRNA mimic (0.25 µg/ml) and performed RNA-seq (Figure 2A). Poly I:C is known to strongly activate OAS and, in turn, RNase L as well as other dsRNA sensing pathways, including PKR, and RIG-I/MDA5. First, we assessed the levels of RNase L activation by measuring the level of rRNA degradation and comparing it to that in cells treated by the direct activator, 2-5A. We found that in all replicates of poly I:C and 2-5A treated cells, the level of rRNA degradation was comparable (within 2-fold), suggesting a similar level of RNase L activation (Figure S1A). Next, we determined the DE genes in WT cells upon poly I:C treatment with DESeq2. We found that poly I:C treatment resulted in a higher number of relatively upregulated DE genes (∼1200, log2 fold changes >1, p_adjusted_ <0.05) as compared to 2-5A treatment (∼700 genes) and that additional pathways were activated, as expected given the broader effects of poly I:C (Figure 2B). In particular, in both WT and *RNASEL* KO cells, PKR and RIG-I/MDA5 pathways were activated, as judged by increased levels of eIF2α phosphorylation (shown previously in ^21^ and discussed further below) and the increase in ISGs (Figure 2C and Table S1), respectively. Consistent with this, we observed that *PPP1R15A* (*GADD34*), a gene involved in the ISR (see below) and known to be transcribed as a result of both interferon production via the IRF3/7 pathway and PKR activation ^34^, also exhibited a relative increase compared to other transcripts (∼32-fold) in poly I:C treated WT cells (Table S1 and Figure 2C).

**Figure 2.**
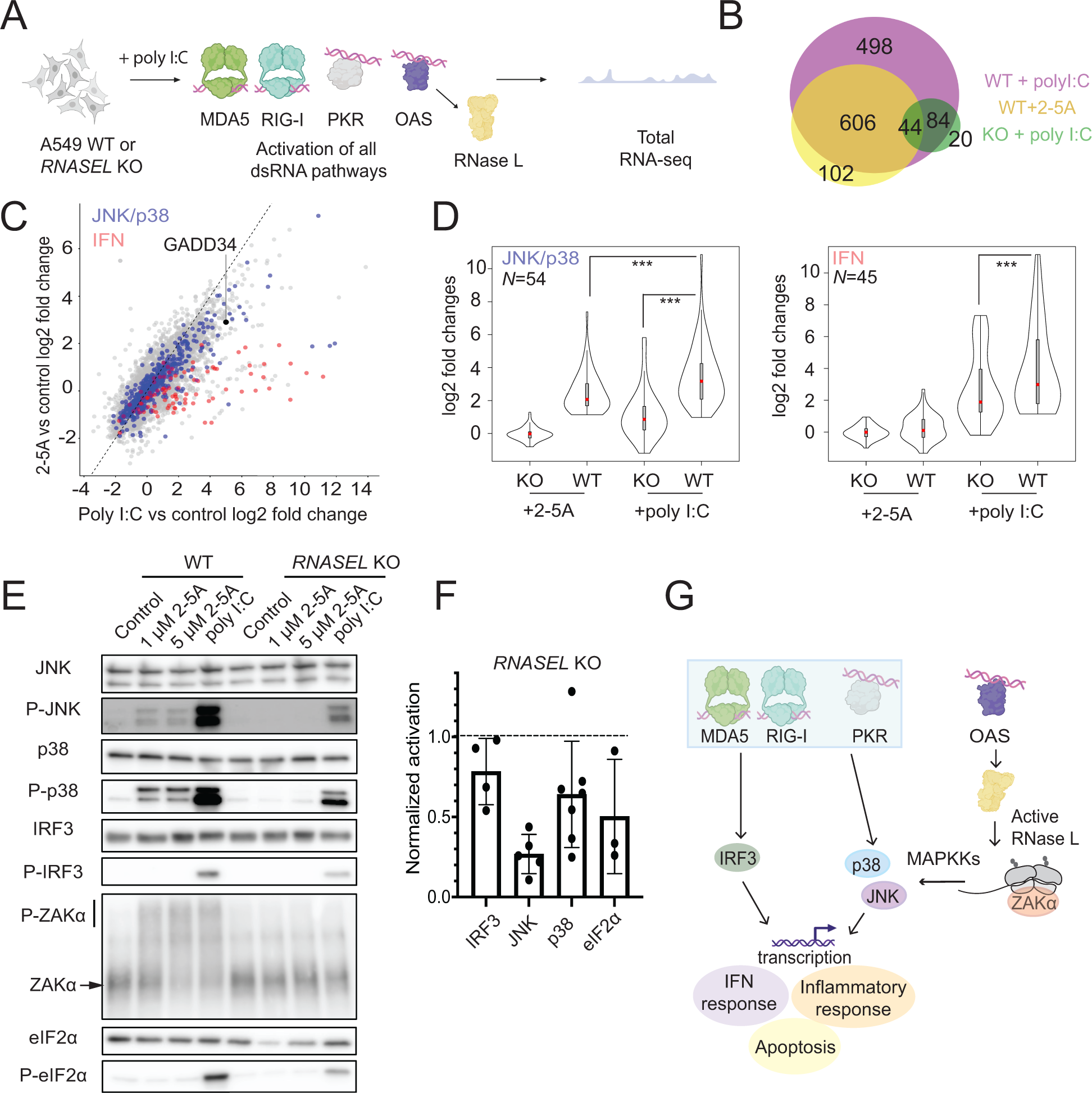
OAS-RNase L and other dsRNA pathways act together to induce the proinflammatory response. **A** Schematics of RNA-seq experiments. A549 cells were transfected with poly I:C (leading to activation of MDA5, RIG-I, and PKR, besides OAS). **B** Venn-diagram showing the overlap of DE genes in cells treated with poly I:C in WT (magenta), 2-5A in WT (yellow) and poly I:C in *RNASEL* KO cells (green, abbreviated to KO). **C** Correlation between differential expression changes in 2-5A (activation of RNase L) vs poly I:C (activation of all dsRNA pathways including RNase L) treated cells. Blue and red dots represent JNK/p38 MAPK activation-related or IFN stimulated genes (as defined in Materials and Methods), differentially expressed in 2-5A or poly I:C treated cells, respectively. Black dot represents *GADD34*. **D** Violin plot representation of genes related to JNK and p38 MAPK activation and interferon stimulation. Statistical testing (paired t-tests) showed significant differences between data sets as indicated (p value <0.001, marked by ***). **E** Effects of activating the OAS-RNase L pathway on phosphorylation of JNK, p38, eIF2α and IRF3 assayed by Western blotting. ZAKα is hyperphosphorylated during dsRNA response. Upper bar shows phosphorylated ZAKα in a Phos-Tag gel (shifted from its expected molecular weight). Lower line points to non-phosphorylated ZAKα. **F** Quantification of western blot for poly I:C treated WT and *RNASEL* KO cells to assess level of activation when RNase L is activated alone or with other dsRNA pathways. Activation levels are normalized to that of poly I:C treated WT cells. Data reveal strong RNase L dependence for JNK and some dependence for p38, eIF2α, and IRF3. **G** Model for RNase L acting synergistically with the other dsRNA pathways to enhance the antiviral effect for JNK and p38 via ZAKα.

Notably, we also found that most genes that are differentially expressed upon 2-5A treatment were also differentially expressed in poly I:C treated cells (Figure 2B, note overlap between yellow- and magenta-colored circles, Pearson’s R^2^=0.92). This suggests that these changes are largely unimpaired in cells treated with poly I:C and are not masked by the effects of the other activated dsRNA pathways. We noted that many genes that did not meet the criteria for significant differential expression in WT cells treated with 2-5A changed in ways that closely mirrored changes observed in cells treated with poly I:C (Pearson’s R^2^=0.75 for DE genes this subset). This correlation indicates that the small relative gene expression changes in 2-5A treated cells that do not meet the DE threshold may indicate partial activation of pathways that get amplified when other dsRNA sensors are triggered in poly I:C treated cells.

To more closely investigate how simultaneous activation of RNase L and other pathways triggered by dsRNA affects the JNK and p38 pathways, we examined the mRNA levels of target genes and phosphorylation of the proteins themselves. Based on our finding above that 2-5A treatment (RNase L activation alone) activates the JNK and p38 pathways (Figure 1E, Figure 2C, blue dots), we defined a subset of 54 transcripts that are commonly associated with JNK and p38 and are activated in 2-5A treated samples (see Methods section for more details). Since ∼50% of the genes downstream of JNK and p38 overlap, we analyzed these MAPK pathways together. We assessed transcriptional induction of JNK/p38 downstream targets under activation of RNase L only (WT + 2-5A), other dsRNA pathways only (*RNASEL* KO cells + poly I:C), or both pathways (WT + poly I:C) (Figure 2D). As expected based on the selection of targets, we found that these JNK/p38-related mRNAs were highly induced in WT + 2-5A treated cells. We also noted some activation due to other dsRNA pathways alone (*RNASEL* KO + poly I:C). Interestingly, when combined, the RNase L and other dsRNA pathways together suggested the effects may be additive since the level of induction was highest in WT cells treated with poly I:C (Figure 2D). We therefore asked whether these trends were supported by the level of JNK and p38 phosphorylation, as measured by western blotting (Figure 2E). In support of additivity between the RNase L and the other dsRNA pathways, we observed the strongest phosphorylation of JNK and p38 when both RNase L and the other dsRNA pathways were triggered compared with either one alone (Figure 2E and F). In addition, we found that ZAKα, which we showed to be phosphorylated by 2-5A treatment (Figure 1E) was also phosphorylated in poly I:C treated WT cells to a similar extent, suggesting that activation of ZAKα by RNase L is not affected by the other dsRNA pathways (Figure 2E). Importantly, ZAKα phosphorylation was not detectable in *RNASEL* KO cells that were treated with poly I:C, suggesting its activation (presumably via ribosome collisions) requires RNase L activity. This model therefore implies that JNK and p38 phosphorylation in the absence of RNase L (Figure 2E) must use a MAP3K other than ZAKα and that, together with ZAKα, affords the strongest induction of p38/JNK targets in cells treated with poly I:C (Figure 1E).

Activation of ZAKα by ribosome collisions can simulataneously occur with activation of the GCN2 kinase, which phosphorylates eIF2α and is also activated by ribosome collisions ^25,27^. Hence we wondered if RNase L could contribute to eIF2α phosphorylation through ribsome collisions. We did not detect eIF2α phosphorylation after 2-5A treatment, consistent with previous reports ^5,21^. However, we found that eIF2α phosphorylation levels were substantially decreased in poly I:C treated *RNASEL* KO cells as compared to WT cells (Figure 2E and F), which is also consistent with previous reports ^17^. This finding shows that activation of RNase L increases eIF2α phosphorylation under some conditions, perhaps by enhancing PKR activation, activating another kinase such as GCN2, and/or reducing the activity of an eIF2α phosphatase.

Next, we assessed if activation of RNase L affects the interferon response that is triggered by dsRNA binding to MDA5 and RIG-I (Figure 2C, red dots). We defined a subset of ISGs (*N* = 45, see Methods for details) that were upregulated in WT cells treated with poly I:C. We then looked at the activation of these genes across our experimental conditions (Figure 2D, right panel). We found no activation of these genes with 2-5A treatment, consistent with previous RNA-seq findings in A549 cells ^16,17,21^, but contrasting with models where RNase L cleaved RNA fragments stimulate other dsRNA sensors, potentially in other cell lines ^18,19^. In cells treated with poly I:C, we observed some decrease in activation for this limited subset of genes in *RNASEL* KO cells compared to

WT cells (Figure 2D). This finding suggests that RNase L may have a minor role in augmenting activation of other dsRNA sensors in these cells ^18,19^. We further investigated these trends by western blotting (Figure 2E and F). In these experiments, we visualized the phosphorylation of IRF3, one of the main transcription factors that induces ISGs in response to interferon. We were unable to detect activation of IRF3 in cells treated with 2-5A but observed activation in cells treated with poly I:C, in agreement with our RNA-seq analysis. We also observed a small decrease (∼20%) in activation of IRF3 in *RNASEL* KO cells treated with poly I:C, in line with our observations from RNA-seq and a previous report ^17^. This supports a model where RNase L may have some role in affecting the response to interferon.

We also asked whether RNase L can contribute to the other pathways that play a role in the dsRNA-induced innate immune response, such as NF-κB and ERK. Activation of NF-κB (p65) was only detectable in poly I:C treated cells and the levels of activation appeared to be similar in both WT and *RNASEL* KO cells, indicating that RNase L has no further effects on this pathway (Figure S2). We also found that 2-5A or poly I:C treatments had no effect on ERK activation in A549 cells, suggesting that the observed effects in our experiments are not due to activation of ERK (Figure S2).

Taken together, our analysis shows that activated RNase L can act together with other dsRNA pathways to promote p38 and JNK signaling, but has only minor effects on the activation of IRF3 (Figure 2G).

### A generic endonuclease, RNase A, induces transcriptional changes similar to those triggered by RNase L

Next, we wanted to assess what changes directly result from the global RNA cleavage that occurs after RNase L activation. RNase L is known to interact with several proteins, including those that make up the cytoskeleton and the nucleopore complex ^35–40^. Thus, it is possible that some of the observed changes upon RNase L activation and dimerization arise from functional protein-protein interactions and not RNA cleavage. To further investigate how global RNA cleavage changes gene expression, we electroporated a generic RNase, RNase A (bovine pancreatic ribonuclease), in *RNASEL* KO A549 cells and performed RNA-seq after 4.5 hours (Figure 3A). RNase A is a small endonuclease that primarily cleaves at pyrimidine bases (U and C) in ssRNAs ^41,42^. As a control, we electroporated similar amounts of Bovine Serum Albumin (BSA) since it is similar in size, but has no RNase activity. We anticipated that RNase A would cause widespread RNA decay but would lack RNase L specific interactions. During our experiments, we followed RNase A activity by monitoring rRNA cleavage, similar to our above experiments with RNase L. We observed that rRNAs are degraded in RNase A electroporated cells, but with different RNA cleavage patterns (Figure S3A). Based on RNase A’s low level of specificity, we expect that most RNA species (including mRNAs) would be targeted and thus fragmented by RNase A.

**Figure 3.**
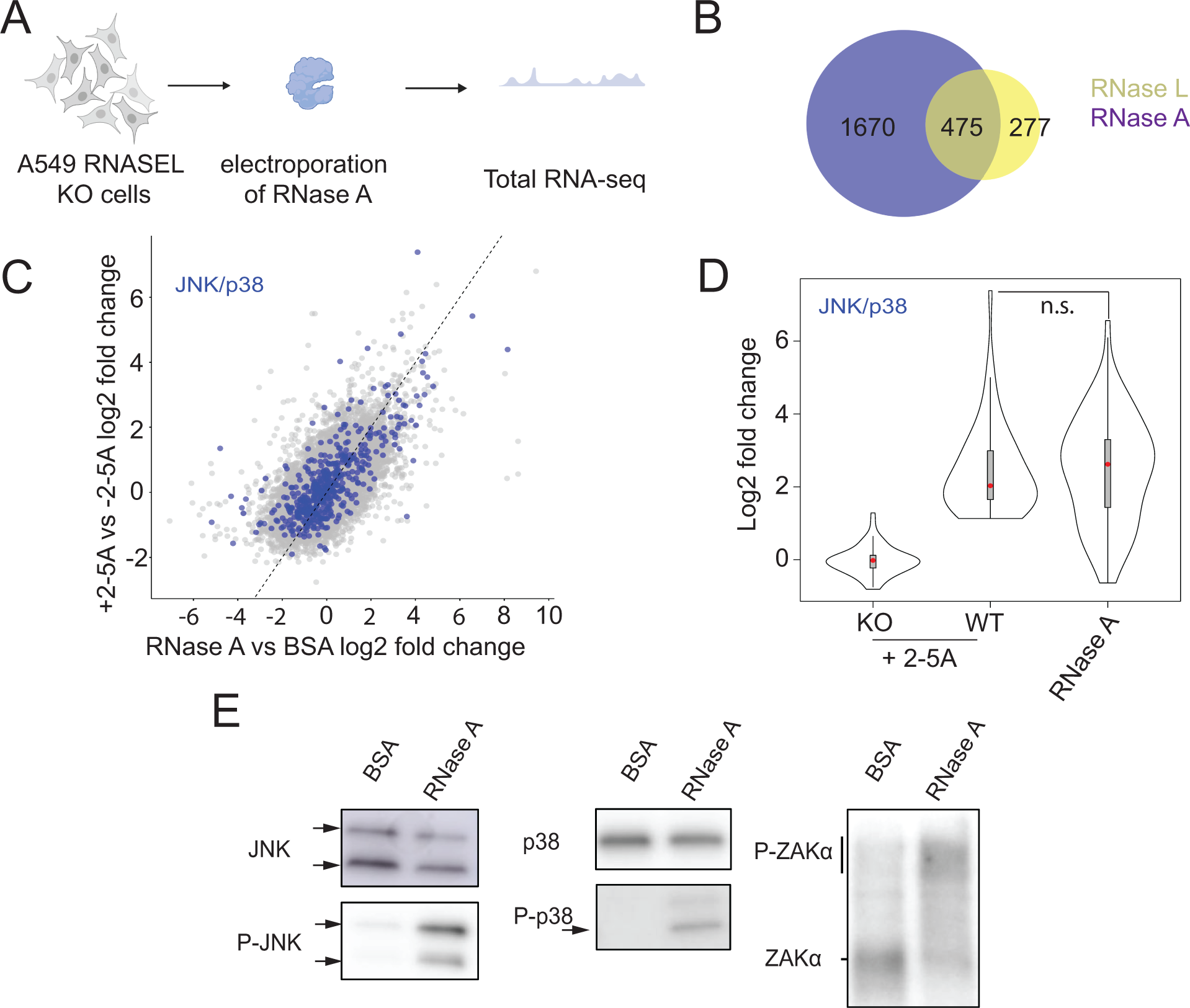
Global RNA cleavage in the cell leads to transcriptional changes. **A** Schematics of RNA-seq experiments. *RNASEL* KO A549 cells were electroporated with RNase A or BSA (control) and RNA-seq experiments were performed as described in Materials and Methods. **B** Venn-diagram showing the number of upregulated genes (defined as p_adjusted_ value <0.05 and log2fold changes >1) in RNase A electroporated samples (purple) and 2-5A treated cells (yellow). **C** Comparison of RNA log2 fold changes between RNase L activated vs RNase A electroporated samples. Blue dots are representing transcripts related to JNK and p38 activation. **D** Violin plot representation of those genes related to JNK and p38 activation (*N* = 54, as defined in Materials and Methods) in RNase A electroporated cells. *RNASEL* KO is abbreviated to KO. Statistical analysis (paired t-test) showed that the log2 fold changes between RNase A electroporated and 2-5A activated samples were not significantly different (p_adjusted_ value >0.05, marked by n.s.). **E** JNK, p38 and ZAKα are activated due to RNase A treatment, as shown by western blots. Different JNK isoforms are marked with black arrows. Upper bar shows phosphorylated ZAKα in a Phos-Tag gel (shifted from its expected molecular weight). Lower line points to non-phosphorylated ZAKα.

We found >1000 differentially expressed genes in RNase A electroporated cells (Table S1). As noted for RNase L activation above, gene expression is reported in absolute terms from the average global loss of mRNA; therefore, genes that appear to be upregulated may simply be less downregulated than those in the same treatment group. Interestingly, most upregulated DE genes found in 2-5A treated cells overlap with those identified here for RNase A activation (∼65 % of those in the RNase L dataset, Figure 3B and C, Figure S3B) and the expression levels showed correlation (Pearson’s R^2^ = 0.39, Figure 3C). This suggests that most RNase L-dependent DE changes are the direct result of RNA cleavage rather than potential indirect non-enzymatic activities.

Next, we assessed if targets of the JNK signaling pathway were upregulated in RNase A electroporated cells as they were in 2-5A treated ones. We found that the JNK and p38 pathway-related mRNAs were upregulated in RNase A electroporated samples (Figure 3D). JNK and p38 phosphorylation were also confirmed by western blotting (Figure 3E). Strikingly, we found that ZAKα was also phosphorylated (Figure 3E). Our results therefore indicate that transcriptional changes, particularly through the JNK and p38 pathways, can be induced by non-specific RNA cleavage, regardless of the effector endonuclease.

### RNA fragmentation by generic RNases induces altORF translation

Previously, we examined the translation that takes place in RNase L activated cells by using ribosome profiling to find where ribosomes translate the reduced pool of mRNAs ^21^. One of the most striking findings from our previous study is that the relative abundance of ribosome footprints increased in non-coding regions of transcripts, including the 5’ and 3’ UTR regions, as well as out-of-frame parts of coding sequences. This phenomenon likely results when ribosomes initiate translation at the first available open reading frame on the 3’ mRNA fragments that are generated by RNase L, a phenomenon that we termed “altORF translation” ^21^. The model predicts that the activity of any generic endonuclease should lead to the same outcome. To test this hypothesis, we electroporated RNase A into *RNASEL* KO A549 cells, as described above, and performed ribosome profiling on those same samples after 4.5 hours (Figure 4A).

**Figure 4.**
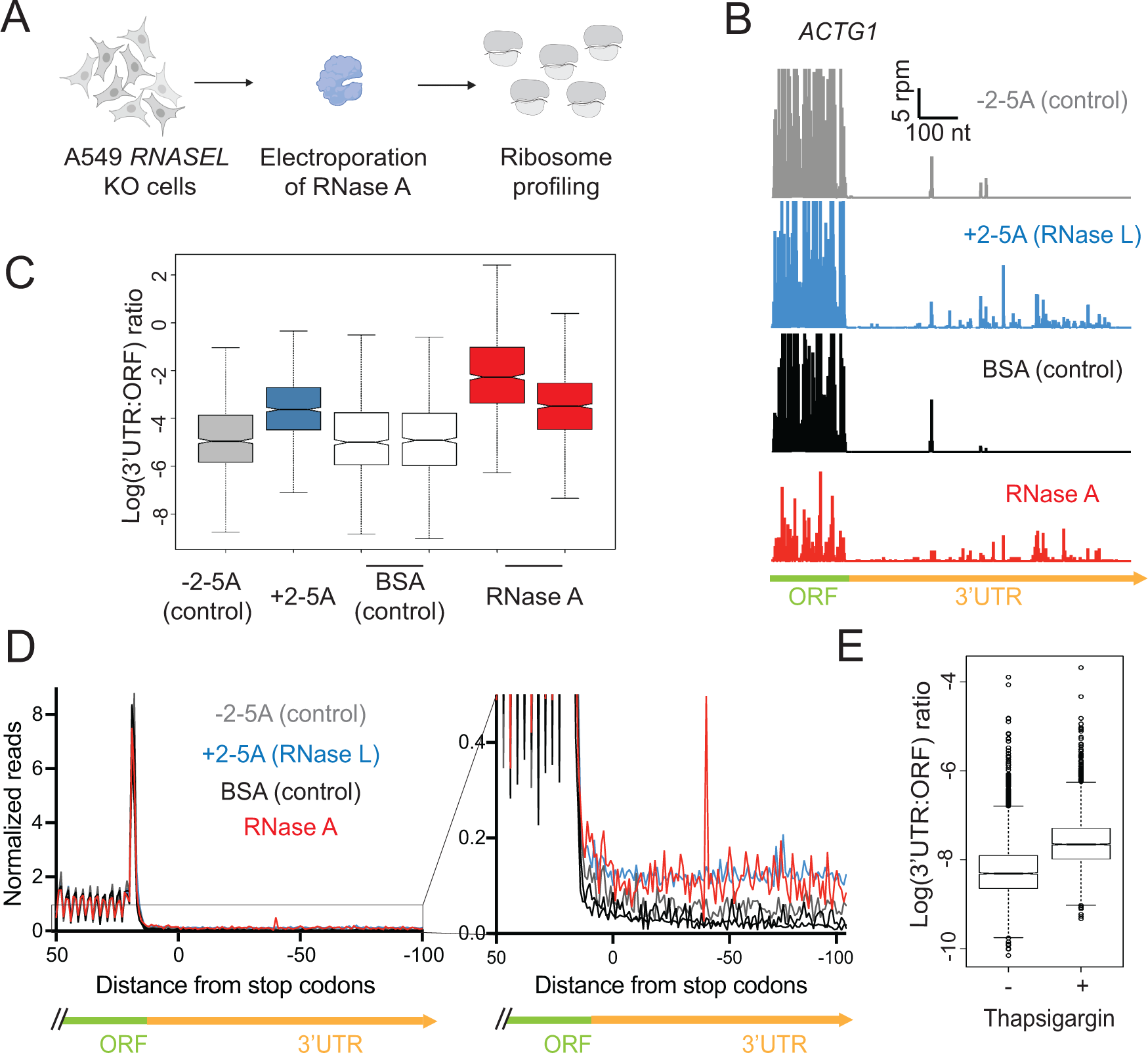
RNA fragmentation leads to alternative ORF (altORF) translation. **A** Schematics of ribosome profiling experiments. *RNASEL* KO A549 cells were electroporated with RNase A or BSA (control) and ribosome profiling experiments were performed as described in Materials and Methods. **B** Ribosome profiling reads in the 3’UTR of the *ACTG1* gene model shows the signature of altORF translation in RNase A electroporated and RNase L activated A549 cells, but not in the controls (-2-5A and BSA). **C** Increased 3’UTR:ORF ratios indicate higher fraction of translation in 3’UTRs when active RNases are present in the cell, but not in controls (-2-5A and BSA). **D** Normalized average ribosome footprint occupancy (metagene plot) around the stop codon of main ORFs (left panel) reveals increased proportion of ribosome footprints in the 3’ UTRs (right panel, zoom in). **E** Increased 3’UTR:ORF ratios indicate altORF translation during thapsigargin treatment and activation of the UPR, but not in untreated cells. In all panels, data shown for RNase L activation (+2-5A) were obtained from ^21^.

The major hallmarks of altORF translation that we previously observed in RNase L activated cells were recapitulated in the cells where RNase A was electroporated (Figure 4B-D). The most striking signature of altORF translation is a relative increase in ribosome footprints in the 3’UTR. This region of the mRNA is rarely translated under normal conditions, but we found ribosomes in this region when RNase L was active ^21^. Cells where RNase A was active exhibited similar ribosome profiling patterns in the 3’UTRs, consistent with the altORF translation model (Figure 4B, *ACTG1* as an example gene model). We also investigated whether this signature was present in RNase-A treated samples globally. To assess characteristics of altORF translation in a global manner, we computed the ratios of ribosome profiling footprint levels in 3’ UTR regions and the main ORFs of mRNAs (3’UTR:ORF ratios) as we did before for active RNase L. We found that in all RNase treated samples, 3’UTR:ORF ratios increased compared to respective controls (Figure 4C) ^21^. Increased 5’UTR:ORF ratios were also observed in RNase L activated cells ^21^ and we found similar patterns when computing 5’UTR:ORF ratios for RNase A treated cells, pointing to higher relative levels of altORF translation in these samples (Figure S4A). Increased reads in the 5’ and 3’UTRs were also observed in metagene averages at the start and stop codon of main ORFs for RNase A electroporated samples as compared to the control (Figure 4D and S4B), further confirming increased relative ribosome occupancy and potential altORF translation in the UTR regions.

The data from cells that were electroporated with RNase A suggest the phenomenon of altORF translation takes place whenever any generic endonuclease, not only RNase L, is active in the cytoplasm. However, electroporation of RNase A is artificial, and we wondered whether another naturally activated endonuclease would trigger the translation of altORFs. The endonuclease IRE1 is homologous to RNase L and plays a pivotal role in the <u>U</u>nfolded <u>P</u>rotein <u>R</u>esponse (UPR) since it is activated by protein misfolding in the ER ^43^. IRE1 is required for cytoplasmic splicing of the *XBP1* mRNA through recognition of specific sequences and structural elements. However, IRE1 can also cleave more generically at UGC sequence motifs. This broader activity is referred to as <u>R</u>egulated IRE1 <u>D</u>ependent <u>D</u>ecay (RIDD) and could potentially lead to altORF translation ^29,44^. To test this hypothesis, we analyzed a publicly available ribosome profiling dataset where human cells were treated with thapsigargin, a potent inducer of the UPR and IRE1 ^45^ (GEO ID: GSE103719). We found that thapsigargin-treated cells also exhibited characteristics of altORF translation, including increased 3’UTR:ORF ratios (Figure 4E). These results show that RNase L is not the only naturally activated endonuclease that can lead to the translation of altORFs.

### RNA fragmentation can modulate translational efficiency

The major effect on gene expression of active RNase L is that the majority of mRNAs are lost, resulting in reduced protein synthesis for most genes. However, transcriptional upregulation counteracts this and, in some cases, overrides (Figure 1D) this effect. In addition, it is conceivable that gene expression from the remaining mRNAs could be tuned at the translational level and thus further refine the amount of protein synthesized from the remaining transcriptome. As an example of this, an increase in the translation of some host mRNAs was reported during vaccinia virus infection, which is a potent activator of RNase L ^46^. In this way, mRNA degradation could have a role in regulating translation.

To investigate whether RNase L activity alters which mRNAs are translated, we took advantage of our ribosome profiling data (reported previously ^21^) and with its matched RNA-seq data (reported here) assessed translational efficiency (TE) of coding sequences. The TE metric is defined as the ribosome footprint level in a coding region, normalized to the corresponding RNA-seq level, and serves as a proxy for the relative loading of a transcript by ribosomes. We therefore plotted the change in ribosome footprint levels against the corresponding change in RNA-seq level (Figure 5A) for genes where either of these values were found to change significantly by DESeq2 (see Methods). Those genes with large changes in TE will appear furthest from the diagonal (slope >1 indicates TE increase; <1 TE decrease). We noted several cases where dots were shifted from the diagonal, suggesting functionally important regulation of translation.

**Figure 5.**
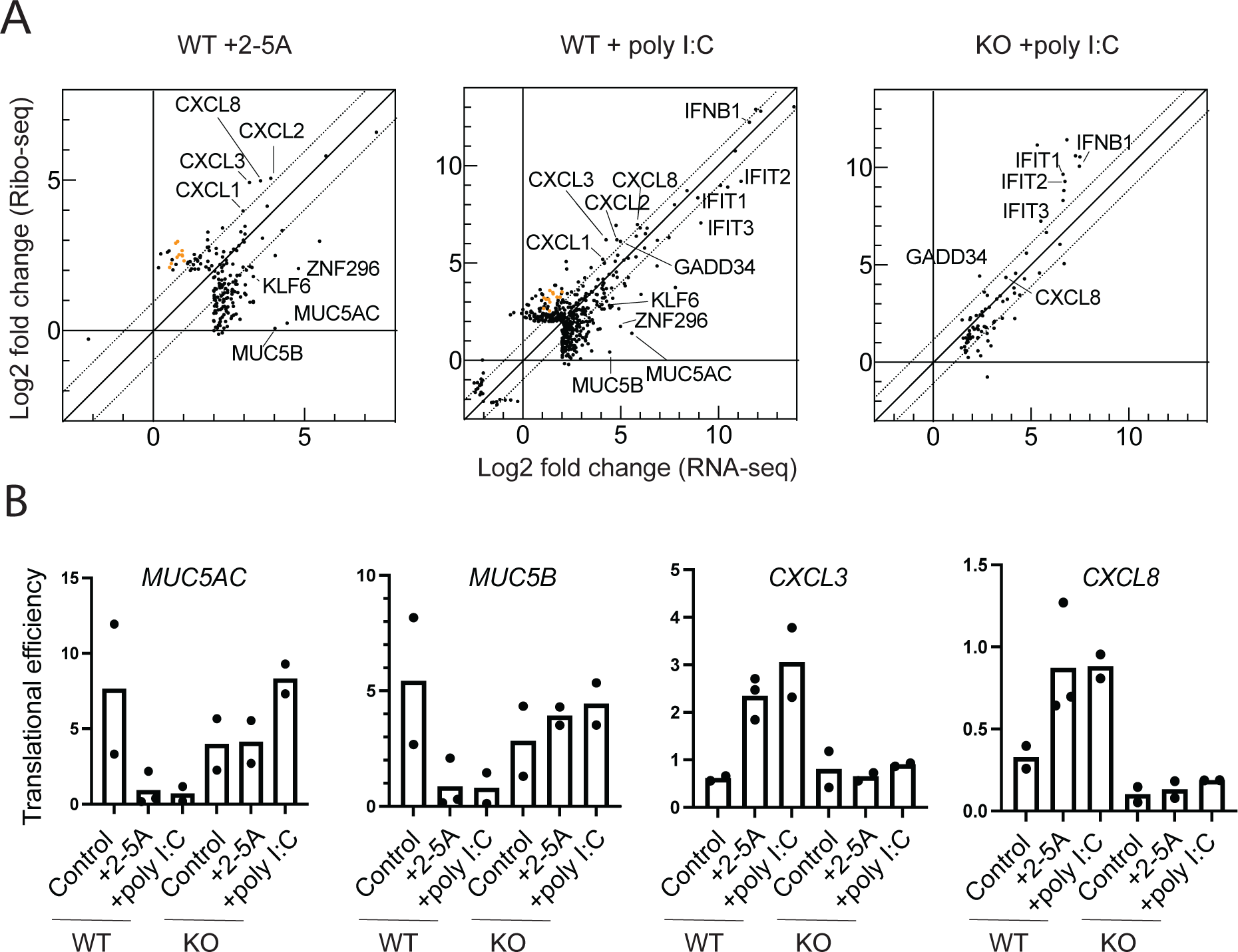
RNase L activation influences translational control and effects of poly I:C treatment. **A**. Comparison of changes in RNA-seq vs ribosome profiling under conditions of 2-5A treated WT cells (left), poly I:C treated WT cells (center), and poly I:C treated *RNASEL* KO A549 cells (right). To emphasize the highest and most significant changes, genes were filtered for the following criteria: log2 fold changes >4 (either RNA-seq or ribosome profiling), p_adjusted_ <0.01, normalized mean counts across the samples >50 (see Materials and Methods for details). Dotted diagonal lines represent TE changes >2 fold. Orange circles represent genes encoded by the mitochondria. **B.** TEs from individual replicates of representative genes with downregulated (*MUC5AC* and *MUC5B*) or upregulated (*CXCL3* or *CXCL8*) translation were computed. TEs calculated as described in Materials and Methods. *RNASEL* KO is abbreviated to KO in all panels.

Some of the largest changes we observed affected a group of transcripts that were expressed at higher relative levels when RNase L was active at the mRNA level (via 2-5A or poly I:C in WT, but not in *RNASEL* KO, cells) but were counterbalanced by downregulation of expression at the translational level (lower TE, Figure 5A, dots to the right of the origin and below the dotted diagonal). This category includes the extracellular matrix proteins *MUC5AC* and *MUC5B* (Figure 5A). We directly computed the TE for these genes for each replicate and found the effect of reduced TE to be consistent (Figure 5B). We also noticed an enrichment for genes encoding proteins involved in transcription or related to transcription in this category (e.g. *KLF6*, *ZNF296*), suggesting that RNase L tunes the production of proteins involved in transcription at the translational level.

In another class of genes, we observed that relative mRNA levels were increased upon RNase L activation (via 2-5A or poly I:C in WT, but not in *RNASEL* KO, cells) and that the level of ribosome footprints also increased to a larger extent and would therefore be expected to augment gene expression (Figure 5A, dots to the right of the origin and above the dotted diagonal). This category included *CXCL1-3* and *8*, genes that encode chemokines and cytokines. We also directly computed the TE for these genes for each replicate and found the effects to be consistent (Figure 5B). Interestingly, we found that mRNAs encoded by the mitochondrial genome also fell into this category (Figure 5A, orange dots). This observation is consistent with a study of TEs during vaccinia infection where higher levels of translation of mitochondrially-encoded transcripts in conjuction of higher oxidative phosphorylation activity was reported. This was proposed to enhance viral replication due to its high energy demand ^46^. Our data therefore suggest that this phenomenon is driven by RNase L.

Intriguingly, we also observed a trend among mRNAs that are induced by dsRNA pathways apart from RNase L, including *IFIT1-3* and *IFNB1* (induced by interferon) and the gene *GADD34* (induced by interferon and the ISR as noted above). These genes exhibited ∼2 fold higher TEs in poly I:C treated *RNASEL* KO cells as compared to WT cells (Figure 5A, dots shift above diagonal in right vs middle panel), suggesting that RNase L may play some role in reducing their expression at the translational level as a way to tune their transcriptional induction.

### RNase L activation modulates the Integrated Stress Response

Given the role of *GADD34* in the ISR, we further examined the ribosome profiling data for clues as to how its translation is regulated by RNase L. The ISR is activated when PKR binds dsRNA and phosphorylates the translation initiation factor eIF2α, ultimately resulting in a loss of its ability to promote translation initiation ^47^. The downstream effects of eIF2α phosphorylation are collectively known as the Integrated <u>S</u>tress <u>R</u>esponse (ISR) ^11,12^. Phosphorylation of eIF2α generally limits the loading of ribosomes on mRNAs in the cell, reducing their TE. However, there is a small class of mRNAs, including *ATF4*, *GADD34* (*PPP1R15A*), and *CHOP* (*DDIT3*), that undergo the opposite response and are translationally activated ^11^. ATF4 and CHOP are transcription factors that induce expression of stress-responsive genes, while GADD34 is a phosphatase of eIF2α and thus serves as a counterbalance to ISR activation ^48,49^. We found that relative transcript levels increased in poly I:C treated WT and *RNASEL* KO cells as compared to respective controls for *GADD34* (as noted above, 5-32-fold) and *CHOP* (5-9-fold) (Table S1). This is consistent with work showing both genes are transcriptionally induced by ATF4 ^50,51^. *GADD34* is likely further induced by the interferon-induction IRF3 pathway ^34^. The *GADD34* mRNA was identified as a strong target of RNase L driven degradation ^17^ and thus absolute levels of *GADD34* are likely lower in WT cells as compared to *RNASEL* KO cells. The transcripts for *ATF4*, *CHOP*, and *GADD34* contain one or more open reading frames in their 5’UTR (uORFs) with unique features that allow them to suppress translation of the main ORF in non-stressed cells, but activate it when eIF2α becomes phosphorylated ^11,52^. As noted above, the TE for *GADD34* was observed to increase in poly(I:C) treated cells (WT or *RNASEL* KO) (Figure 6A), though 2-fold less in WT than RNASE L KO cells.

**Figure 6.**
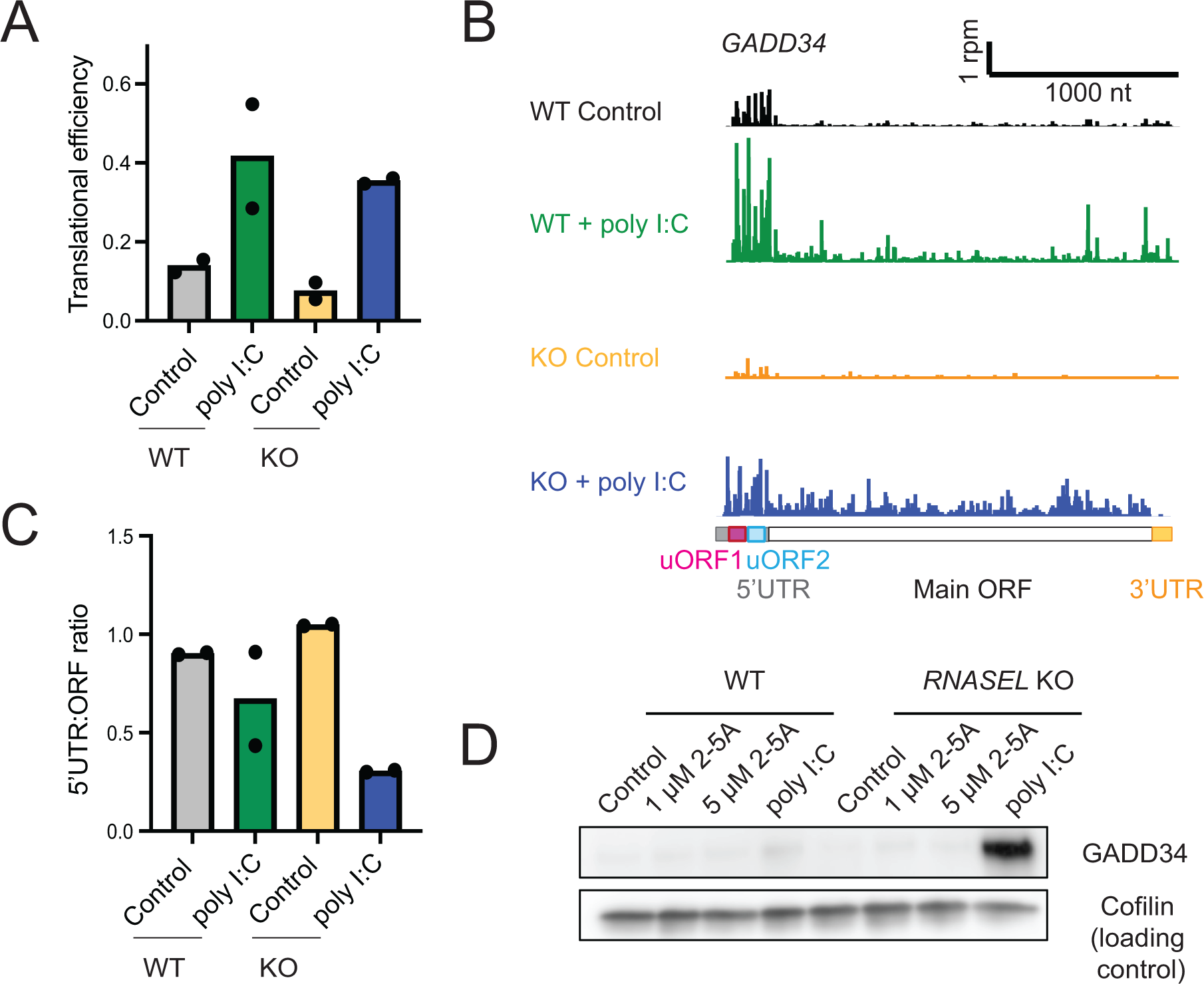
RNase L modulates elements of the Integrated Stress Response. **A** Bar plots representing TEs of *GADD34* mRNA during poly I:C treatment in WT and *RNASEL KO* cells. These data show that TE changes are greater in *RNASEL* KO cells during the ISR as compared to respective controls. **B** Ribosome profiling tracks of gene models for *GADD34* (*PPP1R15A*) in WT and *RNASEL* KO cells during poly I:C treatment suggest a change in 5’UTR vs main ORF translation ratio in *RNASEL* KO cells during the ISR as compared to WT cells and their respective controls. **C** 5’UTR:main ORF ratios of ribosome profiling data in WT and *RNASEL* KO cells with and without poly I:C treatment were computed for individual replicates. Data show this value is more strongly reduced upon poly I:C treatment in cells where RNase L is absent. **D.** Western blot for GADD34 shows increased levels of protein in poly I:C treated *RNASEL* KO cells, reflecting both the lack of mRNA degradation and enhanced TE in cells where RNase L is absent.

Several explanations could account for our observation that the change in *GADD34* TE was less in WT vs *RNASEL* KO cells (Figure 6A). One possibility is that RNase L could directly affect TE by altering the shift in translation from the uORFs to the main ORF. To determine whether RNase L had an effect on uORF-based regulation, we quantified the ratio of ribosome footprints in the 5’UTR and compared it to footprints that mapped to the main ORF (Figure 6B-C, see Methods). We noted a substantial decrease in uORF translation relative to main ORF translation in poly I:C treated WT or *RNASEL* KO cells due to eIF2α phosphorylation, as expected ^52^. However, this decrease was less in the case of the WT cells compared to *RNASEL* KO cells (Figure 6B-C), suggesting that RNase L activation may have some role in affecting this shift. Similar RNase L dependent shifts in 5’UTR vs main ORF ribosome footprints for *ATF4* and *CHOP* suggest this regulation may also extend to those genes (Figure S5A-B). It has been shown that the *GADD34* mRNA is heavily targeted by RNase L, making the mRNA and protein product in poly I:C treated *RNASEL* KO cells much more detectable than in WT cells ^17^. Consistent with this, we also found that GADD34 protein levels increased in poly I:C-treated *RNASEL* KO cells (Figure 6D). Thus, this mechanism where RNase L suppresses main ORF translation augments the underlying trend in expression that is driven by mRNA decay.

Taken together, these results show a role of RNase L in inhibiting the translation of *GADD34*, thus modulating the ISR during the innate immune response, and suggests the RNase L may have a broader role in regulating translational control.

### Translation of IFIT mRNAs is inhibited by activation of RNase L

Since we observed that poly I:C induced higher translation efficiency in *RNASEL* KO vs WT cells for the genes *IFIT1*, *2*, and *3* (Figure 5A, right panel), we asked whether this trend could be explained by uORF-mediated translational control as in the case of the ISR-related mRNAs. IFIT proteins are one of the key antiviral proteins that are stimulated by interferon and inhibit viral translation ^53^. Since these genes have low expression under conditions without poly I:C stimulation, we focused on data from cells treated with poly I:C.

To determine whether uORFs were translated on these mRNAs, we examined ribosome densities in their 5’UTR in ribosome profiling data. We found that 5’UTRs of *IFIT1*, *2* and *3* contained ribosome peaks at start codons of potential uORFs (Figure 7A and Figure S6A, green lines). The uORFs used near-cognate start codons (CUG or GUG instead of AUG, Figure 7A and Figure S6A). These data suggest a model for translational control where the uORFs inhibit main ORF translation and it is conceivable that this inhibition could be regulated by eIF2α phosphorylation. As in the case of *GADD34* and other genes, this apparent uORF inhibition may be regulated by RNase L since there was lower 5’UTR translation relative to main ORF translation in poly I:C treated *RNASEL* KO vs WT cells that was correlated with higher TE (Figure 7B and C). These data further imply that expression of these genes may be controlled by inhibitory uORFs and that this mechanism may depend on RNase L (Figure 7D). We also found that IFIT1 and IFIT2 protein levels increased in poly I:C-treated *RNASEL* KO cells compared to WT cells (Figure S6B). This change is consistent with the reduction in translation efficiency or mRNA loss in WT cells.

**Figure 7.**
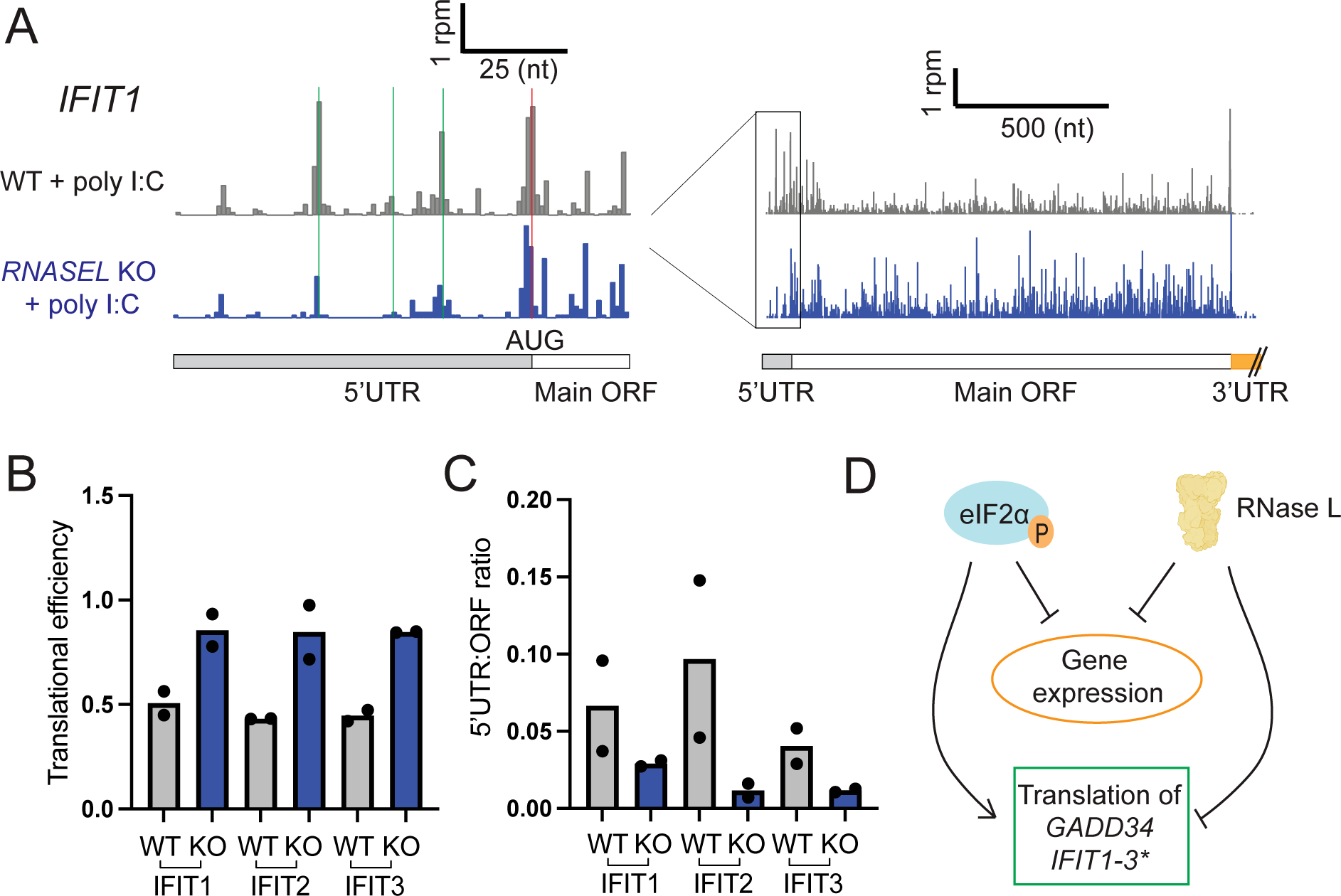
IFIT mRNAs undergo changes in translation that depend on RNase L. **A** uORFs were identified on *IFIT1* mRNAs based on the increase in ribosomes density at alternative start codons (CUG, green markings) in poly I:C treated cells. Red marking shows the start codon of the main ORF. **B** TEs of *IFIT1-3* mRNAs are decreased in RNase L deficient cells (comparing WT and *RNASEL* KO poly I:C treated cells) consistently across replicates. **C** *IFIT1-3* mRNAs were found to have less 5’UTR relative to main ORF translation in poly I:C treated *RNASEL* KO (abbreviated to KO) cells as compared to WT cells. **D** Schematics of how eIF2α phosphorylation and activation of RNase L affects gene expression in the cell. While both mechanisms lead to reduced gene expression (less translation when eIF2α is phosphorylated and less overall mRNA when RNase L is activated), eIF2α phosphorylation increases translation of *GADD34* but RNase L reduces its translation. Similarly, RNase L reduces *IFIT* mRNA translation. It may also be increased via uORF regulation under conditions of eIF2α phosphorylation (marked by asterisk since further confirmation is needed).

## Discussion

In this study, we examined how direct RNase L activation by 2-5A, and attendant global RNA degradation in the cell, affects gene expression. Our work shows that the global loss in mRNA is compensated by increases in transcription, particularly via activation of ZAKα, and tuned by changes in translation.

We showed that one way mRNA cleavage leads to new transcription is via activation of the RSR. RNase L activation alone (via 2-5A) turns on the p38 and JNK MAPK pathways, independently of other dsRNA sensors that are activated by poly I:C (Figure 1). While JNK activation was previously documented in several cell lines, simultaneous p38 activation was not observed when RNase L alone was activated ^14,23^. Since poly I:C treatment strongly induces p38, it is possible that weak activation of p38 was masked in previous studies, or the extent of activation could be cell line dependent. The activation of JNK is known to be associated with the RSR ^54,55^ and it was initially suggested to be involved in innate immunity due to damage to rRNA by RNase L ^22,23^. However, more recent studies showed that even with their rRNA damaged by RNase L, ribosomes can still translate ^16^ and may therefore not be neccesarily responsible for RSR activation. Recently, it was found that the RSR can be activated by the ribosome collision sensor and MAP3K, ZAKα ^25,27^. We established that ZAKα can also be triggered directly by RNase L activity, but not by the other dsRNA pathways. Moreover, it is likely that other mechanisms, beyond the RSR, exist to activate transcription in response to global RNA degradation. For instance, global RNA loss in the cytoplasm was proposed to free up RNA binding proteins that, in turn, modulate transcription due to their relocation to the nucleus ^56^.

Our observation raises the question of why ribosome collisions might be occurring in these cells. Ribosome collisions are known to form when a ribosome stalls during elongation and upstream ribosomes bump into it ^57^. One possibility is that mRNAs undergoing active translation are cleaved by RNase L, resulting in the ribosomes stalling whenever they reach a 3’ end of a 5’ cleavage fragment. Prior work using ribosome profiling demonstrated this kind of 3’-end stalling and formation of ribosome collisions when the homologous nuclease, Ire1, cleaves mRNAs during the unfolded protein response in fission yeast ^29^. Consistent with the idea of an increase in ribosome collisions during viral infection, it was shown that vaccinia virus infection leads to an increase in ubiquitination of the ribosomal protein uS10 by the ubiquitin ligase ZNF598, which is known to be directly activated by ribosome collisions ^58^. Alternatively, ribosome collisions could form for other reasons, including the widespread translation of alternative (non-canonical) ORFs that we previously reported ^21^, since such ORFs may be less evolutionarily tuned to minimize slow translation ^59^. We also note that ZAKα has been reported to be somewhat activated by individual stalled ribosomes, implying ribosome collisions are not strictly required for the effects observed here ^60^. It is also possible that ZAKα could be activated in other ways that involve the cleaved RNA fragments through a yet unknown mechanism. The effect of RNase L-induced activation of the RSR was previously shown to be JNK-mediated programmed cell death in response to viral infections ^22^. Interestingly, both ZAKα and RNase L activation were shown to trigger the inflammasome independently ^20,28^. Considering our data, it is plausible that RNase L also triggers inflammasome activation through ZAKα, and contributes to cell death.

Previous RNA-seq studies hinted that RNase L activation may act together with the other dsRNA pathways to amplify the innate immune response ^17^. Our findings here confirm this synergy by showing that activation of JNK and p38 occur by both RNase L dependent and independent routes. Since impairment of RNase L activation can have serious health consequences linked to a reduced ability to fight viral infections ^61,62^, it is possible that the loss of this two-pronged activation weakens other branches of the immune response that are independently triggered by dsRNA. In addition, our observation that ERK is not activated and p38 is activated by 2-5A has not been consistently observed in previous studies ^14,23^, further suggesting that the activation of kinases by RNase activity is regulated by other factors, possibly in a cell-line dependent way.

Interestingly, our data also suggest that RNase L may lead to further regulation of eIF2α kinase activity. Our analysis of eIF2α phosphorylation levels in cells treated with poly I:C shows that phosphorylation is higher when RNase L is active. This is consistent with previous reports showing that besides PKR, another kinase can phosphorylate eIF2α during poly I:C treatment and it is dependent on RNase L activity ^17^. A possible explanation is that RNase L activates the eIF2α kinase GCN2, which is sensitive to ribosome collisions ^25,63–65^ and enhanced by ZAKα activation ^25^. Intriguingly, our analysis and previous studies ^5,21^ of eIF2α phosphorylation levels suggest direct RNase L activation via 2-5A does not activate any kinase of eIF2α. Therefore, enhanced phosphorylation of eIF2α when RNase L is active likely relies on additional factors that are activated by poly I:C but are absent when only RNase L is active.

Beyond control of new transcription, we also examined roles for RNase L in changing which transcripts are translated. We noted several prominent cases (Figure 5) where mRNA TEs were changed when RNase L was activated by 2-5A. The mechanism underlying these translational changes remains unclear, however, several possible explanations exist. Certain transcripts, such as *IFNB1*, are retained in the nucleus via direct interactions with nucleopores when RNase L is activated ^35^ or a viral nuclease is activated ^66^. This mechanism would explain the apparent decrease in translational efficiency of this gene (decreased TE in poly I:C treated WT as compared to *RNASEL* KO, Figure 5) and possibly other transcripts due to selective retention in the nucleus. In addition, TE changes could arise because the availability of ribosomal subunits and the corresponding translation machinery, such as initiation factors, likely increase because the overall level of mRNA is greatly reduced upon RNase L activation ^16,67^. Such a dramatic change in this “ribosome homeostasis” (as defined, ^68,69^), or balance between mRNA and the translational machinery, has been shown to result in changes in which transcripts are favored for translation ^70,71^. Such a result is thought to occur because mRNAs that are generally less efficient at recruiting ribosomes can be more heavily translated due to an abundance of ribosomal subunits promoting translation via mass action.

We also focused on changes in translation efficiency related to the ISR since it is a well-established regulatory pathway known to rely on control of translation via phosphorylation of eIF2α by the dsRNA sensor PKR ^11,52^. One consequence of eIF2α phosphorylation is the translational activation of a set of genes, such as *GADD34*, due to a shift in translation initiation from uORFs to the main ORF ^11^. Intriguingly, we found that activation of RNase L overrides this effect of eIF2α phosphorylation and limits the translational activation of *GADD34*. This translational suppression would serve to further enhance the inhibition of this gene due to its known susceptibility to RNase L mediated degradation (Figure 6) ^17^. One possible explanation for the inhibitory effect of RNase L on the translation of this gene is that the availability of eIF2α may increase when RNase L is active due to the general reduction in demand for translation, as noted above. As a result, the impact of phosphorylation of eIF2α on translation initiation may be lessened, thus preventing the shift from uORF to main ORF translation on these transcripts. Alternatively, since RNase L cleaves rRNAs, it is plausible that this damage could alter the choice of start codon (uORF vs main ORF) by scanning 40S subunits. Another possibility is that we previously showed ^21^ that RNase L activation leads to an increase in the proportion of ribosome footprints in 5’UTRs globally and this holds true for *GADD34* (Figure 6B-C). This effect may arise from direct initiation on mRNA cleavage fragments generated when RNase L cleaves in the 5’UTR. This change in initiation could therefore lead to the reduced translational activation of *GADD34*. Interestingly, we also found that eIF2α phosphorylation levels were decreased in poly I:C treated *RNASEL* KO cells compared to WT. Since GADD34 directly dephosphorylates eIF2α, we postulate that increased amounts of GADD34 protein in poly I:C treated *RNASEL* KO cells could contribute to the decrease of eIF2α phosphorylation.

Additionally, we found that RNase L-dependent changes in translation efficiency of *IFIT1*, *IFIT2*, and *IFIT3* (as well as other interferon-related transcripts) is correlated with an apparent shift in uORF to main ORF translation. One potential explanation for this effect is that these *IFIT* genes, much like *GADD34*, could be activated by the ISR. In this scenario, RNase L would similarly serve to limit their activation. While the expression of these genes is likely already downregulated at the mRNA level by RNase L mediated degradation, this mechanism offers additional control and suppression. Such dampening could be important for limiting autoimmune conditions. For example, impairments in the OAS-RNase L pathway is one of the contributing factors for MIS-C (<u>M</u>ultisystem Inflammatory <u>S</u>yndrome in <u>C</u>hildren) ^72^. In the case of MIS-C, the disease appears to manifest due to the lack of RNase L’s control of inflammation. Thus, active RNase L may serve a dual role in innate immunity, activating key pathways while counterbalancing others.

Finally, we found that the generic endonuclease RNase A evokes a transcriptional response, including that mediated by ZAKα, that is very similar to that of RNase L (Figure 3). In addition, RNase A, or the ISR-mediated endonuclease IRE1, can also trigger the translation of altORFs, much like RNase L (Figure 4). This evidence suggests other endonucleases, such as the viral endonuclease Nsp1 or the apoptotic host endonuclease EndoU ^66,73,74^, could potentially cause similar outcomes to RNase L. It is intriguing that virally-encoded endonucleases may bring about outcomes that are similar to host endonucleases and suggests the effects of generic endonuclease activation can be tuned to benefit either the virus or the host. For example, in the case of Nsp1, nuclear mRNA retention and cytoplasmic mRNA degradation were shown to be important for limiting the interferon response and promoting viral replication ^35,66^. On the other hand, more limited dampening of these pathways, particularly for *IFNB1* ^17^ and *IFIT* genes shown here (Figure 7), could be important for preventing autoimmune conditions or cytokine storms, such as noted above. Further studies are needed to elucidate these possibilities.

Overall, our data offer a model where activation of RNase L induces ribosome stalling and likely collisions, leading eventually to cell death via key transcriptional pathways (Figure 1) related to the kinase ZAKα. This suggests that ZAKα could be a potential target for therapeutics aimed at modulating RNase L’s effects. This is especially relevant since RNases are already being used or tested for cancer therapy, such as RNase A ^75,76^. We speculate that a component of the mechansim behind these RNase therapies could result from ZAKα activation via ribosome collisions that occur as a result of RNase activity in the cytoplasm. Thus, modulating the levels of ZAKα activation could be the key for safety and efficacy of these treatments. In addition, RNase L may have other roles in affecting the ISR via translation of ISR target genes. The effects of RNase L on transcription and translation may be important for reinforcing or dampening effects evoked by other dsRNA sensors, pointing to linkages between pathways in the innate immune response. Further study is needed to dissect these connections that bring together the entire spectrum of mRNA activities, including synthesis, translation, and degradation.

## Funding

This research was supported by the Intramural Research Program of the NIH, The National Institute of Diabetes and Digestive and Kidney Diseases (NIDDK) (DK075132 to N.R.G.) and the Postdoctoral Research Associate Training Program (PRAT) at the National Institute of General Medical Sciences (NIGMS) (FI2GM128743 to A.K.).

## Acknowledgements

We are thankful for the insightful discussions with Drs. Bret Hassel, Alan Hinnebusch, Jon Lorsch, Stacy Horner and Tom Dever. A549 cells were a kind gift from Dr. Bernie Moss. We also thank Dr. Behdad Afzali and Dr. Daniel Chauss for their assistance with electroporation experiments and the assistance of the genomic cores at the National Institute of Diabetes and Digestive and Kidney Diseases (NIDDK) and the National Heart, Lung, and Blood Institute (NHLBI). A.K. is grateful to have been chosen for a MOSAIC K99/R00 award (K99 GM143484) and for associated mentoring support from the American Society of Cell Biology partnership activities.

## Data availability

All raw sequencing (fastq) and processed alignment (wig) files are available at NCBI GEO in record GSE244176 (ribosome profiling) and GSE244125 (RNA-seq). It can be accessed with the following token for reviewer access: gdcbakkitjgrzkh (GSE244176) and anypgeyefjoxzkj (GSE244125). Custom code used for ribosome profiling in this study is available on Github: https://github.com/guydoshlab. Gene expression analysis workflow is available on https://github.com/TriLab-bioinf/TK_65_RNAseL_manuscript.

## Author contributions

A.K. designed and performed experiments, analyzed the data and wrote the paper. H.A.L. performed RNA-sequencing data analysis, contributed to DESeq analysis, and edited the manuscript. A.V.D. performed western blotting experiments. N.R.G. designed experiments, developed software, and wrote the paper.

## Materials and Methods

### Cell culture

WT and *RNASE* L KO A549 lung carcinoma cells were cultured in RPMI (Gibco Cat # 60870127) complemented with 10% Fetal Bovine Serum (Gibco FBS). A549 cells were tested and negative for Mycoplasma contamination throughout the study. Mycoplasma testing was performed using eMyco Valid Mycoplasma PCR detection kit (BioLink) or ATCC Universal Mycoplasma Detection Kit. Cells were incubated at 37°C in the presence of 5% CO_2_. Cell lines used in this study were a kind gift of Dr. Bernie Moss (NIH) and generated as described in ^77^.

### rRNA cleavage assay

Total RNA was extracted from ∼3×10^5^ cells using Direct-zol mini kit (Zymo) or Trizol reagent (Invitrogen) according to the manufacturer’s protocol. The amount of total RNA was computed by absorbance at 260 nm measured by NanoDrop (Thermo Fischer Scientific) and then diluted to 50-500 ng/µl. Then RNA samples were run on a Bioanalyzer 2100 (Agilent) using the RNA 6000 Nano kit (Agilent) or TapeStation with the Agilent RNA ScreenTape assay. Minute run-to-run shifts in band sizes are due to inherent limitations of the instrument.

### Sample preparation for RNA-seq and ribosome profiling

Cells used in experiments were grown until they reached near confluency (70-90% confluent) in a T75 flask. Then cells were transfected with the pre-incubated mixture of 1 µM 2-5A or 0.25 µg/ml poly I:C (Invitrogen, low molecular weight) and Lipofectamine 3000 (7.5 µl/6-well) in Opti-Mem serum free media as recommended by the manufacturer for 4.5 hours. Control cells (definied as untreated or -2-5A) were transfected with lipofectamine only. 2-5A was synthesized as described before ^21^. For electroporation of RNase A or BSA, confluent T75s (∼6-8×10^6^ cells) were trypsinized and resuspended in MaxCyte electroporation buffer and electroporated in 3XOC25 cuvettes (MaxCyte) in the presence of 160 ng/ml RNase A or BSA with standard settings for A549 in the MaxCyte ATX electroporator. Cells were plated on T75 flasks and collected after 4.5 hours incubation. Then cell lysates were prepared and collected as for ribosome profiling ^21^. The same samples were split for RNA-seq and ribosome profiling.

### Ribosome profiling

Ribosome profiling of RNase A and BSA electroporated cells was carried out as previously reported ^21^, except that rRNA depletion was carried out by siTools riboPool rRNA depletion kit designed for human ribosome profiling. The ribosome profiling data was processed and signatures of altORF translation were assessed as before ^21^. Sequencing was performed on HiSeq 3000 (single end 50) at the NHLBI or NIDDK DNA Sequencing and Genomics Core. These ribosome profiling data and previously published data were used for analysis of translation efficiency (see below).

### Total RNA-seq

Total RNA was extracted from 100 µl of lysates prepared for ribosome profiling using Direct-zol mini kit (Zymo) or Trizol reagent (Invitrogen) according to the manufacturer’s protocol. The amount of total RNA was computed by absorbance at 260 nm measured by NanoDrop (Thermo Fischer Scientific) and then diluted to 50-500 ng/µl. Then RNA samples were run on Bioanalyzer 2100 (Agilent) using the RNA 6000 Nano kit (Agilent). Samples for sequencing were subject to rRNA depletion and library preparation by Illumina Truseq kit. Sequencing was performed on HiSeq 3000 or Novaseq 6000 (paired end 50 or 75 cycle) at the NHLBI DNA Sequencing and Genomics Core.

## Computational analysis

### Processing and alignment of RNA-seq reads

After removing residual adapters from sequencing reads with the bbduk.sh tool from bbtools (v37.62), sequencing data were mapped to a database of highly abundant rRNA and tRNA sequences with bowtie2 (v 2.4.5) ^78^. Unmapped reads were then mapped to the human reference genome (GRCh38) with STAR (v2.7.10a) using default parameters except for alignSJDBoverhangMin=1, outFilterMismatchNmax=2, alignEndsType=EndToEnd, and leaving multimaper reads mapping to less than 11 loci ^79^. Duplicated reads were removed with picard MarkDuplicates (v2.26.10) ^80^ and quantification of fragments per gene was carried out with the featureCounts tool from Subread software (v2.0.1) ^81^ using Ensembl gene annotations (Ensembl version hg38) by counting read-pairs falling within coding regions (CDS) as fragments and adding fractional counts for multimapper reads. BigWig files were generated with bamCoverage tool from DeepTools (v3.5.1) ^82^.

### Analysis of RNA-seq data

Processed bulk RNaseq data was analyzed in R using the following workflow: https://github.com/TriLab-bioinf/TK_65_RNAseL_manuscript. Differential gene expression analysis was performed with the R package DESeq2 (v1.34.0) ^31^. GO enrichment analysis were carried out with the gseGO function from the R package ClusterProfiler (v4.2.2) ^83^.

To identify genes associated with JNK/p38 and interferon in Figures 2C and 3C, we utilized gene collections from Harmonizome 3.0 ^84^ that were based on Gene References Into Function (GeneRIFs, JNK, p38 datasets, Table S3). Within these datasets, we used a subgroup of genes that were upregulated with 2-5A treatment (JNK/p38 plots, *N*=54, Table S3) to create violin plots in Figure 2D and Figure 3D. For interferon-related genes in Figure 2, we used the hallmark datataset from the Molecular Signature Database ^85^ (“interfron-gamma-response,” Table S3) for a subgroup of genes that were upregulated in poly I:C treated WT cells (interferon-related plots, *N*=45, Table S3). We performed paired t-test to determine significant differences between the groups within each plot (see legends for more details). To perform correlation analysis between datasets from cells treated with 2-5A and poly I:C, we performed a sequential filtering process to compare different subsets of genes according to differential expression status (see R workflow).

### Processing of ribosome profiling footprint data

Ribosome profiling data was processed as described before to remove linkers and duplicates ^21^. After processing, reads were 5’ aligned with STAR (v2.8.2a) to the human genome (hg38, UCSC). Multimapping was allowed up to 200 times (-- outFilterMultimapNmax option) and the maximum mismatch/read was set to 1 (-- outFilterMismatchNmax option). For both the RNA-seq and ribosome profiling data CDS read counts were generated by featureCounts tool from Subread software (v2.0.1) including multimappers as fractional counts and reads on exon-exon junctions (-M -- fraction and -J options, respectively).

### Analysis of translational efficiencies

We used our DESeq2 pipeline as described above for RNA-seq to obtain differential expression values (log2 fold changes) for the ribosome profiling experiments. These changes (RNA-seq and ribosome profiling log2 fold change values) were plotted against each other in Figure 5A to visualize where translation efficiency (TE) was changing. We used a filter for log2 fold changes >4 and p adjusted value <0.01 as the definition for points we considered for TE changes for all datasets. We further filtered these sets for normalized mean counts across the samples >50 to eliminate low expressing genes. Of this reduced dataset, we excepted TE changes >2 (defined by dotted lines on Figure 5A) for further consideration.The output of the DESeq2 analysis for the RNA-seq and ribosome profiling are provided as Supplemental Table 1 and 2.

For further analysis of individual genes, we computed the TE directly by dividing reads per million (rpm) values for ribosome profiling and RNA-seq data that mapped to the coding sequence. This allowed us to directly compare replicates within a treatment group.

### Analysis of translation for altORF analysis and individual genes

For more detailed analysis of ribosome profiling data, we implemented a reduced-transcriptome alignment and python analysis pipeline as described elsewhere ^21,86^. We used the Ref-Seq Select+MANE (ncbiRefSeqSelect), as downloaded from UCSC on 14 April 2020, for this alignment after removal of duplicates on alt chromosomes. Using our python tools (writegene2) to output reads for individual gene tracks, we then computed the ratio of rpm values that mapped to the UTR to those that mapped to the main ORF (Figure 6, 7 and S5), based on the defined annotations. We also used these tools to create metagene plots (Figure 4 and S4) using the (metagene function). When uORFs overlapped the main ORF for *ATF4*, the overlap region was assigned to the UTR for the calculation of UTR to ORF ratio. In general, we assume a shift of 12 nt between the 5’ end of a read and the P site of the ribosome. Data are displayed in individual tracks to correspond with ribosome P sites.

### Western blotting

Protein levels were assessed by SDS-PAGE electrophoresis and immunoblotting during 2-5A and poly I:C treatment. For western blotting cells were grown on a 6-well plate up to ∼70% confluency then transfected with 2-5A or poly I:C as described above. Cells were trypsinized after 4.5 hours (Gibco TrypleExpress) and collected by centrifugation at 200 g for 5 minutes. The cell pellet then was resuspended in 90 µl of 10 mM Tris-HCL pH=8.0, 150 mM NaCl and 1 % triton-X and incubated on ice for 5 minutes. Samples were then centrifuged for 1500 rpm, 4 minutes at 4°C. The supernatant was then mixed with equal amount of 2X SDS-PAGE buffer (Quality Biological) and samples were sonicated with a Brenson sonicator for 10-15 seconds until the sample was no longer viscous. 5-10 µl of sample was run on a 4-20% gradient Mini-PROTEAN Tris-HCl gel (BioRad) or 7% PhosTag (Wako) gel prepared according to manufacturer’s instruction and as described before ^25^. Proteins were transferred to a 0.2 µm PVDF membrane using the Trans-Blot Turbo System (BioRad, 1.3 A, 7 minutes for 4-20% gradient Mini-PROTEAN Tris-HCl gel or 1.3 A for 14 minutes for PhosTag gel). Membranes were blocked in EveryBlot Blocking Buffer (BioRad) for 10 minutes at room temperature. This was followed by incubation with primary antibodies overnight in TBST (Tris-buffered saline plus 0.1% tween) and 1% milk. The following antibodies were used: anti-p38 MAPK (Cell signaling Technologies #9212, 1:2000 dilution), anti-P-p38 MAPK (Cell signaling Technologies, 1:2000 dilution, #9211), anti-IRF3 (Cell signaling Technologies #11904), anti-P-IRF3 Ser396 (Cell signaling Technologies #4947), anti-ZAK (Bethyl A301-993A), anti-P-JNK (Thr183/Tyr185) (Cell signaling Technologies #4668), anti-JNK (Cell signaling Technologies #9252), anti-p65 (Cell signaling Technologies #8242), anti-ERK1/2 (Cell signaling Technologies #4695), anti P-ERK1/2 (Cell signaling Technologies #4370), anti-IFIT1 (abcam #236256), anti-GADD34 (abcam #236256), anti-eIF2α (Bethyl, #A300-721A), anti-P-eIF2α (abcam #32157). After washing 3 times for 5 minutes in TBST, secondary antibodies were incubated with the PVDF membrane for 1 hour at room temperature (Goat anti-rabbit antibody, BioRad, 1:3000). Then, after additional washes (3 times, 5 minutes each) in TBST, the PVDF membrane were incubated with Clarity Western ECL substrate (BioRad) for 5 minutes and proteins were visualized by Amersham Imager 600. Western blots were repeated for 2-4 times using biological replicates and blots were quantified using Fiji ^87^. Replicate images will be provided on Mendeley Data.

## Figure Legends

**S1 Related to Figure 1.**
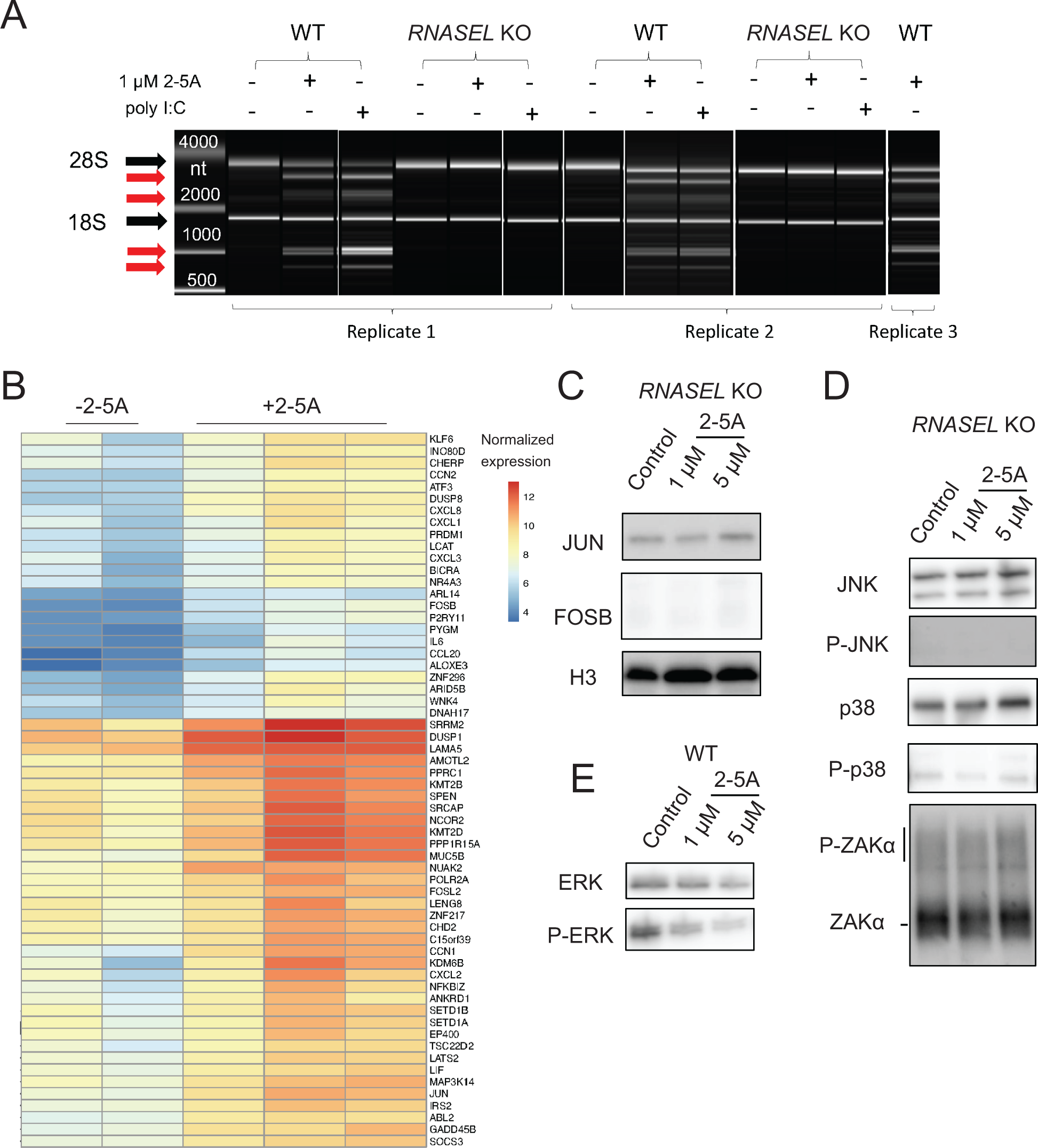
**A** Cleavage patterns of total RNA extracts show activation of RNase L in 2-5A or poly I:C treated WT, but not *RNASEL* KO, A549 cells. rRNA cleavage assay performed on a BioAnalyzer. Black arrows indicate the 28 and 18S rRNAs and red arrows show RNase L degradation products. **B** Heatmap representation of the most highly expressed genes during RNase L activation. Trends are consistent across replicates. **C** Western blots show that transcription factors FOSB and JUN do not increase in *RNASEL* KO cell due to 2-5A activation. **D** Western blots show JNK, p38 and ZAKα are not activated in *RNASEL* KO cells after 2-5A treatment. **E** Western blots show ERK is not activated in WT cells after 2-5A treatment.

**S2 Related to Figure 2.**
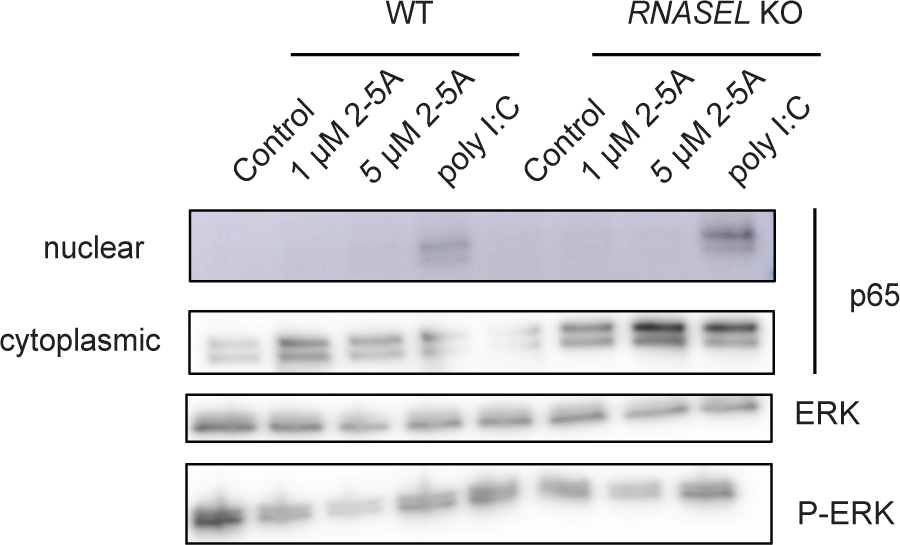
Western blots show no detectable effect of RNase L activation on NF-κB activation by poly I:C and no detectable activation by 2-5A. ERK activation was not observed during 2-5A or poly I:C treatment.

**S3 Related to Figure 3.**
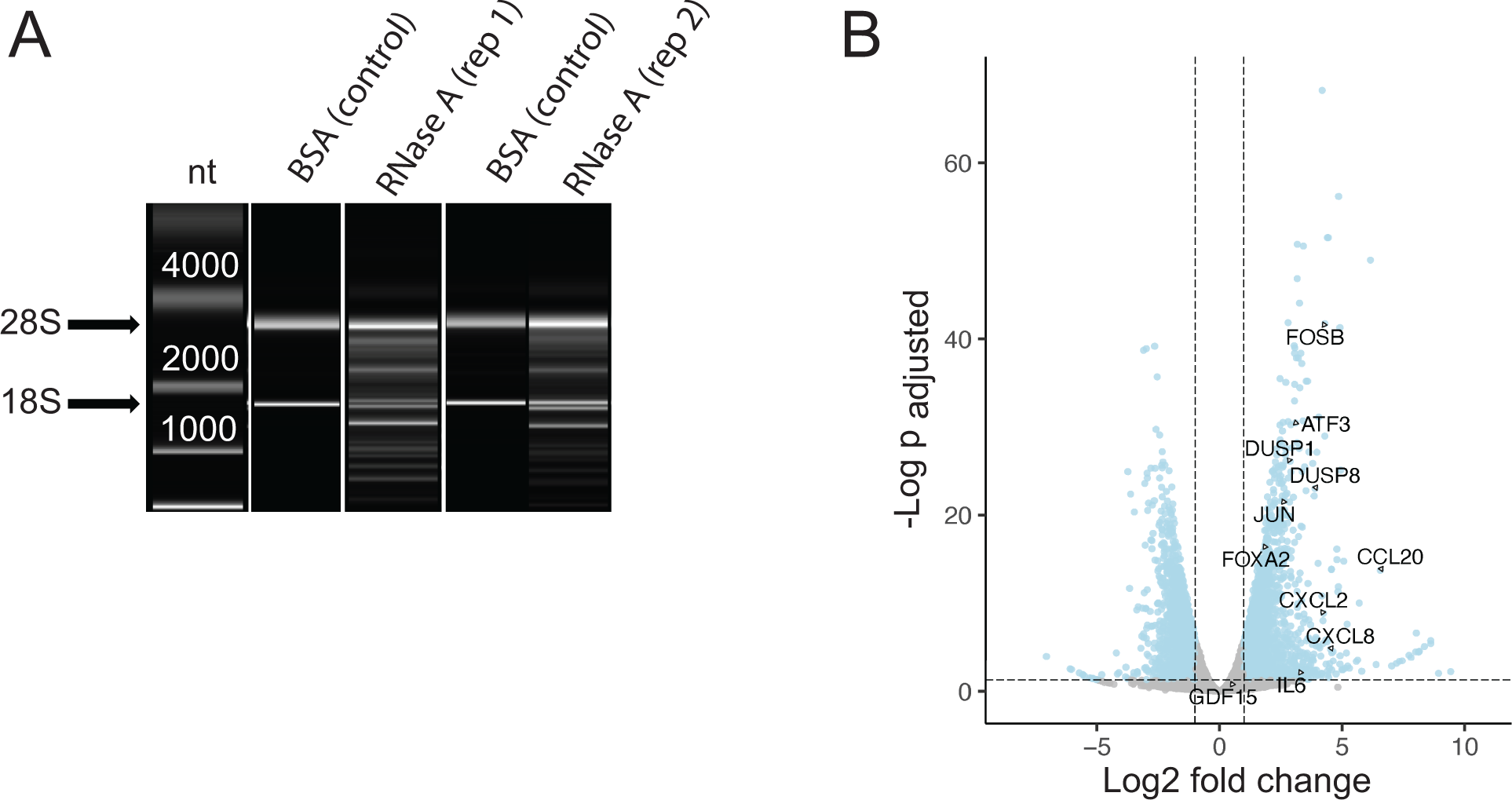
**A** Cleavage of RNAs in the cell occurs when RNase A is electroporated into cells, but not when BSA is electroporated, as observed by rRNA cleavage assays performed on BioAnalyzer. Arrows indicate the 28S and 18S rRNAs. **B** Volcano plot showing upregulation of example proinflammatory cytokines and transcription factors during RNase A electroporation. Differentially expressed genes define as p_adjusted_ value <0.05, log_2_fold change >1.

**S4 Related to Figure 4.**
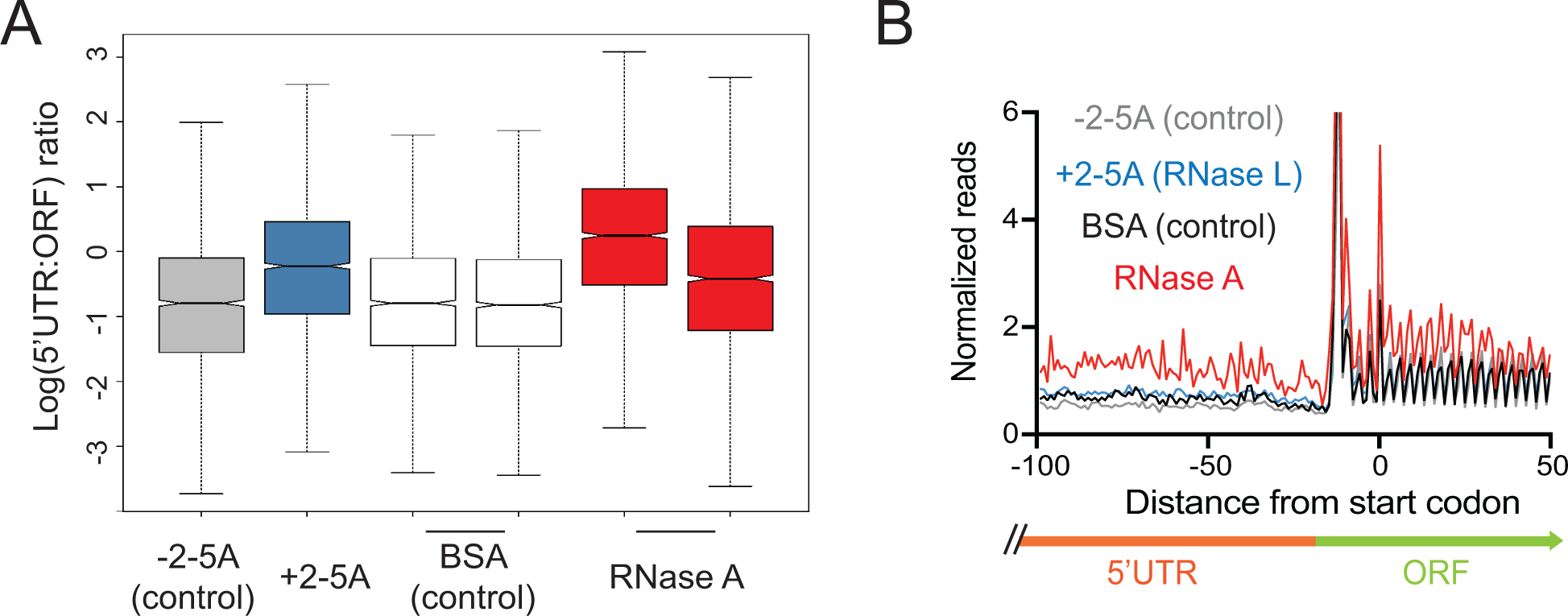
**A** Increased 5’UTR:ORF ratios indicating higher relative uORF translation when active RNase is present in the cell, but not in controls (-2-5A and BSA electroporated). **B** Normalized average ribosome footprint occupancy (metagene plot) around the start codon of main ORFs reveals increased relative ribosome footprint levels in the 5’ UTRs when an active RNase is present vs the respective control. In all panels data shown for RNase L activation (+2-5A) was obtained from ^21^.

**S5 Related to Figure 6.**
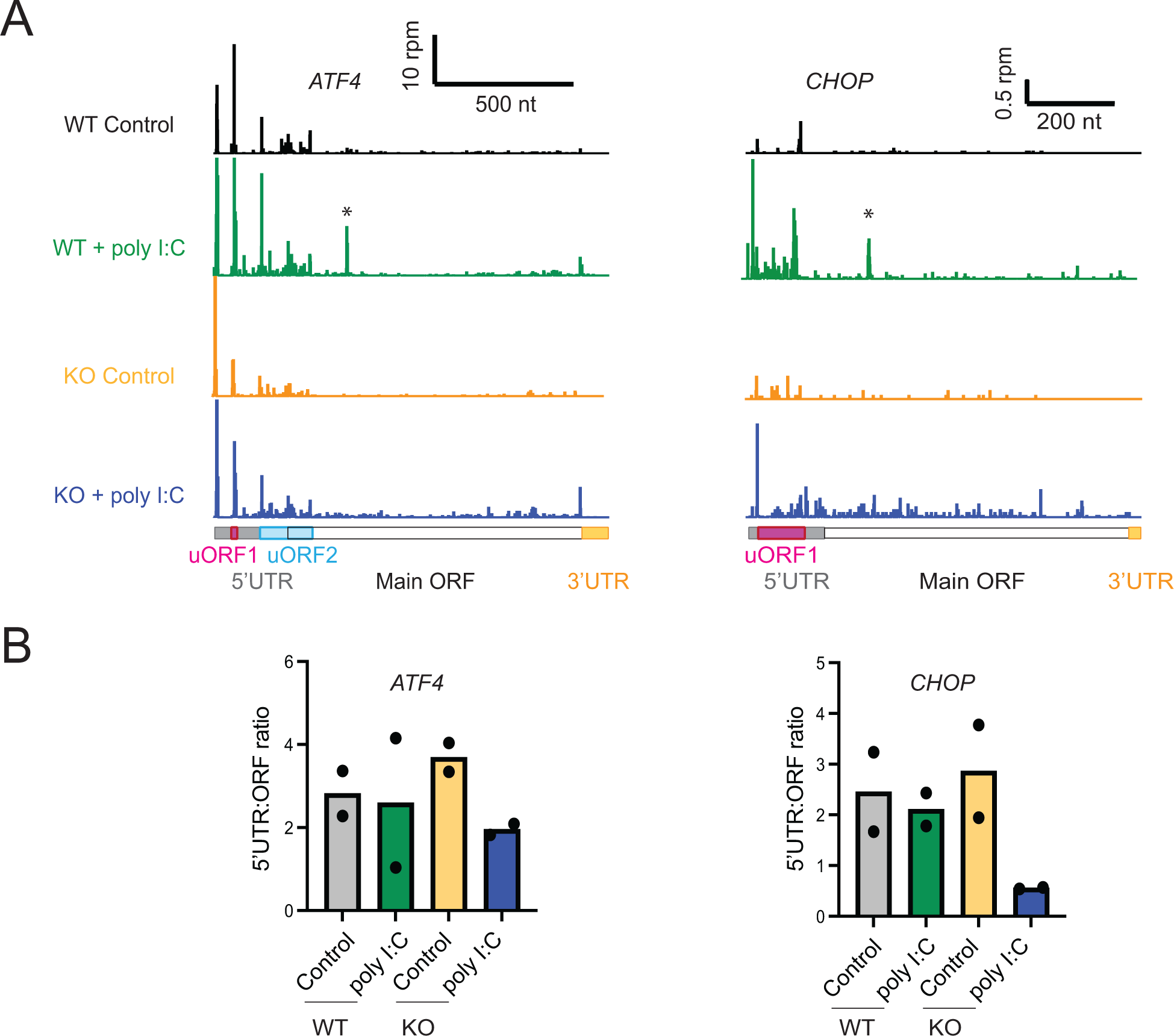
**A** Ribosome profiling tracks for gene model of *ATF4* and *CHOP* in WT and *RNASEL* KO cells during 2-5A or poly I:C treatment. Data show the poly I:C dependent shift toward main ORF vs 5’UTR translation is greater when RNase L is absent. Asterisks show RNase L dependent ribosome profiling peaks in 2-5A and poly I:C treated cells that likely correspond to altORF translation initiation events. **B** 5’UTR:main ORF ratios computed from ribosome profiling data in WT and *RNASEL* KO cells during poly I:C treatment for *ATF4* and *CHOP*.

**S6 Related to Figure 7.**
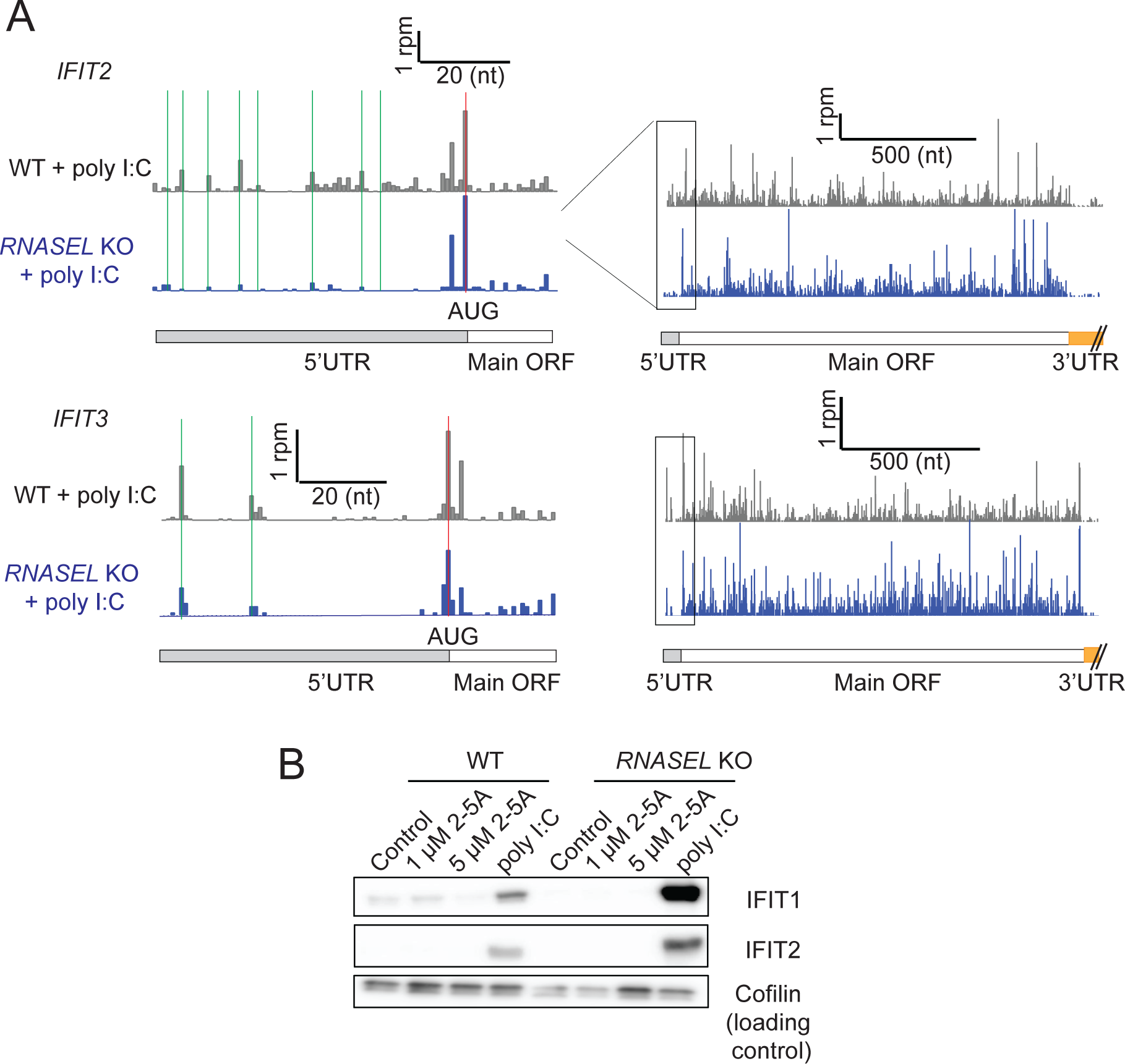
**A** uORFs were identified on *IFIT2* and *3* mRNAs based on the increase in ribosome densities on alternative start codons (CUG or GUG, green markings) in poly I:C treated WT or *RNASEL* KO cells. Red marking shows the start codon for the main ORF. **B** Western blot for *IFIT1* and *IFIT2* shows increased protein levels in poly I:C treated RNase L KO cells, suggesting control of TE or RNA degradation reduce levels in WT cells

**Table S1.** Results tables from DESeq2 analysis for RNA-seq (WT and *RNASEL* KO, +/- 2-5A, +/- poly I:C, BSA vs RNase A electroporated).

**Table S2.** Results tables from DESeq2 analysis for ribosome profiling (WT and *RNASEL* KO, +/-2-5A, +/- poly I:C).

**Table S3.** List of genes that are upregulated by JNK/p38 in WT 2-5A treated cells and interferon response in WT poly I:C treated cells. This list was used to create violin plots in Figures 2D and 3D. In addition, a longer list of all genes related to JNK/p38 or interferon are given, as derived from the Harmonizome and Hallmark datasets, respectively (see Methods), used in Figures 2C and 3C.

## Notes

### Competing Interest Statement

The authors have declared no competing interest.

### Summary of Updates

Minor revisions throughout manuscript and new data in Figure 2.

